# Genetic dissection of MutL complexes in Arabidopsis meiosis

**DOI:** 10.1101/2024.09.28.615604

**Authors:** Nadia Kbiri, Nadia Fernández-Jiménez, Wojciech Dziegielewski, Esperanza Sáez-Zárate, Alexandre Pelé, Ana Mata-Villanueva, Juan L. Santos, Mónica Pradillo, Piotr A. Ziolkowski

## Abstract

During meiosis, homologous chromosomes exchange genetic material through crossing-over. The main crossover pathway relies on ZMM proteins, including ZIP4 and HEI10, and is typically resolved by the MLH1/MLH3 heterodimer, MutLγ. Our analysis of plant fertility and bivalent formation revealed that the MUS81 endonuclease can partially compensate for the MutLγ loss. Comparing genome-wide crossover maps of the *mlh1* mutant with ZMM-deficient mutants and lines with varying HEI10 levels reveals that while crossover interference persists in *mlh1*, it is weakened. Additionally, *mlh1* show reduced crossover assurance, leading to a higher incidence of aneuploidy in offspring. This is likely due to MUS81 resolving intermediates without the crossover bias seen in MutLγ. Comparing *mlh1 mlh3 mus81* and *zip4 mus81* mutants suggests that additional crossover pathways emerge in the absence of both MutLγ and MUS81. The loss of MutLγ can also be suppressed by eliminating the FANCM helicase. Elevated expression of *MLH1* or *MLH3* increases crossover frequency, while their overexpression significantly reduces crossover numbers and plant fertility, highlighting the importance for tight control of MLH1/MLH3 levels. By contrast, PMS1, a component of the MutLα endonuclease, appears not to be involved in crossing-over. Together, these findings demonstrate the unique role of MutLγ in ZMM-dependent crossover regulation.

## INTRODUCTION

Meiosis is a reductive cell division that results in the formation of gametes or spores, thereby enabling sexual reproduction (1). It plays a vital role in generating genetic diversity within populations (2). Several mechanisms operate during meiosis to foster the creation of new genetic variations. One such mechanism is crossover recombination, where fragments are reciprocally exchanged between homologous chromosomes. At least one crossover per chromosome pair is necessary for the accurate segregation of chromosomes during meiosis, a phenomenon known as crossover assurance. Failure to generate crossovers can lead to the formation of unbalanced spores, ultimately reducing fertility and potentially causing aneuploidy in the offspring (3).

In plants, as in many eukaryotes, multiple crossover pathways exist, with the dominant one being the Class I crossover pathway, driven by a group of proteins collectively known as ZMM. ZMM proteins are essential for creating an environment that stabilizes recombination intermediates and facilitates their resolution into crossovers. In *Arabidopsis thaliana*, the ZMM pathway accounts for at least 85% of all crossovers, so the loss of any ZMM component results in a drastic reduction in both crossover events and fertility. In budding yeast, most intermediates stabilized by ZMM proteins result in crossovers, but in many species, like Arabidopsis, the number of ZMM-bound intermediates is about 20 times higher than the final number of crossovers (4). This pathway is also characterized by crossover interference, ensuring that crossovers are more widely spaced along the chromosome than would be expected by random distribution. This regulation of crossover placement is achieved through the ZMM proteins ZYP1 and HEI10. In Arabidopsis, the *zyp1* loss of function mutation eliminates crossover interference (5, 6). Furthermore, the expression level of HEI10 directly affects crossover frequency and the strength of interference (7, 8). The number of HEI10 foci gradually decreases during prophase I through a process called coarsening, which ultimately determines crossover positions along chromosomes (8–10).

The MutLγ heterodimer, composed of the MLH1 and MLH3 proteins, is regarded as the primary endonuclease responsible for crossover formation within the ZMM pathway. Unlike MUS81 and other known endonucleases, which produce crossovers and non-crossovers at a 1:1 ratio, MutLγ exhibits a strong preference for resolving recombination intermediates as crossovers. Recent studies in yeast have elucidated the mechanism behind this bias (11, 12). This unique property of MutLγ ensures that recombination intermediates designated for crossover formation by the ZMM pathway are almost exclusively resolved as crossovers, allowing for precise regulation of crossover numbers necessary to maintain crossover assurance. However, MutLγ complex is typically not classified as an integral component of the ZMM pathway because the mutant phenotypes differ considerably from those of other *zmm* mutants. The number and distribution of MLH1 foci during pachytene are correlated with late recombination nodules and the frequency of Class I crossovers (13–15). Nevertheless, Arabidopsis *mlh3* mutants exhibit only a 60% reduction in crossover numbers, as opposed to the 85% reduction typically observed in *zmm* mutants and show a 25-hour delay in prophase I, displaying univalents at metaphase I (13). *MLH1* is expressed in both somatic and reproductive tissues, while *MLH3* expression is largely confined to flower buds (15–17). Furthermore, both *mlh1* and *mlh3* mutants show partial infertility (13–15).

In addition to MLH1 and MLH3, PMS1 is another MutL homolog that has been identified in Arabidopsis. While the MLH1/MLH3 heterodimer plays a key role in crossover formation, the MLH1/PMS1 heterodimer (MutLα) has been proposed to be involved in the correction of different classes of DNA mismatches (18). In yeast *pms1* mutants, the frequency of crossovers remains unchanged, but post-meiotic segregation issues arise due to errors in repairing heteroduplexes formed during homologous recombination (19). In *A. thaliana*, the absence of *PMS1* results in increased somatic ectopic recombination and reduced fertility (20). However, the meiotic phenotype of Arabidopsis *pms1* mutants has not yet been analyzed.

Class II crossovers, which are insensitive to interference, constitute up to 15% of the total crossovers in Arabidopsis and are primarily dependent on MUS81 (21, 22). Additionally, a second non-interfering crossover pathway has been identified in this species, which depends on FANCD2 (23, 24). Furthermore, there are anti-recombinase pathways that specifically limit MUS81-dependent crossovers. In Arabidopsis, the most effective components of these pathways involve DNA helicases FANCM and RECQ4. Mutants deficient in the genes encoding these proteins exhibit an increase in Class II crossovers and a loss of crossover interference (25–28).

In this work, we demonstrate that the function of MLH1 and MLH3 in crossover formation and distribution is significantly different from the role of ZMM proteins. Unlike in *zmm* mutants, crossover interference and crossover assurance are still preserved in *mlh1 mlh3* double mutants, though weakened. MUS81 can repair recombination intermediates protected by the ZMM pathway, but its unbiased resolution of these intermediates prevents it from fully compensating for the loss of MutLγ. Moreover, our results suggest that the loss of the MLH1/MLH3 heterodimer activates additional pathways, beyond MUS81, in the formation of crossovers. In contrast, PMS1 does not seem to be directly involved in crossover formation.

## METHODS

### Plant material

Arabidopsis seeds were sown on hydrated soil and stratified in the dark for 48h at 4°C. They were then cultivated in growth chambers under a 16h day/ 8h night photoperiod, 150 μmol light intensity, 21°C day and night, and 70% humidity. Col-0 (N1092), L*er*-0 (NW20), *hei10-2* (SALK_014624), *mlh1-2* (GK-067E10), *mlh3-1* (SALK_015849), *mlh3-2* (SALK_067953), *pms1-3* (SALK_124014), *fancm-9* (SALK_120621), *zip4-2* (SALK_068052), *mus81-1* (GK-113F11), *mus81-2* (SALK_107515), were purchased from Nottingham Arabidopsis Stock Centre (NASC). *mlh1-3* (SK25975) was shared by Raphaël Mercier. *mlh1-1* and *pms1-1* (T-DNA original mutants) were shared by François Belzile. The fluorescent tagged line (FTL) Col-*420* was shared by Avraham Levy (29). The different mutants were genotyped using the primers listed in Supplementary Table S1.

### Fertility assays

The seed set was quantified using five siliques starting from the seventh oldest silique of the main stem. The collected siliques were discolored in 96% ethanol and pictured using the Zeiss Lumar V12 Fluorescence Stereomicroscope at the magnification 6.4×, then processed using ImageJ. Pollen viability was investigated as previously described (30, 31). About 500 pollen grains from three replicates (∼1500 events total) were processed for each genotype. The mounted samples were observed using the Leica DM4 B at magnification 20×.

### Seed scoring

Local meiotic crossover recombination frequency is quantified using the seed-based Fluorescent Tagged Line (FTL) (32) and the CellProfiler *SeedScoring* pipeline (33). The frequency of segregation of the two fluorescent cassettes, eGFP and dsRed present at known positions, corresponds to the recombination frequency of the tested line. The *SeedScoring* pipeline recognizes single seed objects and attributes an intensity of fluorescence. The identified objects are categorized as non-color or colored seeds. Recombination Frequency (RF) in centimorgan (cM) is calculated as follows: RF = 100 × (1 – [1 – 2(N_G_ + N_R_) / N_T_]1/2), where N_G_ is green-only fluorescent seeds, N_R_ is red-only fluorescent seeds and N_T_ is the total number of seeds.

### Cytology techniques

Chromosome spreading was performed from at least three replicates per genotype as described in (34). The number of univalents, bivalents, and chiasmata were quantified from DAPI-stained pollen mother cells at metaphase I using a Leica DM4 B epifluorescence microscope equipped with a Leica DMC5400 20-megapixel color CMOS camera, previously described (35). Fluorescence in situ hybridization (FISH) was performed as described by Sanchez-Moran *et al.* (2001) (36), with minor modifications, on specimens where flower bud fixation and male meiocyte spreading were carried out according to (37). The DNA probes used were 45S rDNA, pTa71 (34) and 5S rDNA, pCT4.2 (39). Metaphase I images were scored to determine chiasma frequency and bivalent configurations (ring/rod) per chromosome. The cells were imaged using an Olympus BX61 epifluorescence microscope equipped with an Olympus DP70 digital camera.

### CRISPR-Cas9 mutagenesi

was used to generate null *mlh1* mutants in Col and L*er*, as described in (40). Three sgRNAs were targeted to the region from the 4^th^ intron to the 6^th^ exon of *MLH1*, targeting both splicing variants (Supplementary Figure S2 and Table S2). Four independent mutants were selected and sequenced. Col background mutants showed the same 462 bp genomic deletion, and RT-seq showed a 298 bp deletion at the transcript level, introducing multiple stop codons and a frameshift. L*er* background mutants showed a single nucleotide insertion in the 5^th^ exon, introducing a frameshift and multiple stop codons.

### Lines overexpressing *MutL* genes

were generated by independently introducing extra copies of *MLH1*, *MLH3,* or *PMS1*. Genomic sequences of the different genes were cloned with their respective endogenous promoters or in frame with the meiosis-specific *DMC1* promoter. Cloning primers are listed in Supplementary Table S3. Wildtype plants were transformed using *A. tumefaciens* floral dipping, and transformants were selected using BASTA.

### Genome-wide crossover mapping by F_2_ sequencing

Whole genome sequencing libraries were constructed based on the protocol described in (35, 41–43). gDNA was CTAB-extracted from 288 Col/L*er* F_1_ plants rosette leaves, then isolated using chloroform, precipitated with isopropanol, and purified via ethanol precipitation. The obtained gDNA was suspended in TE and its quality and concentration were checked. The samples were diluted to 5ng/μL and tagmented with an in-house produced Tn5 transposase loaded with the Tn5ME-A (5′-TCGTCGGCAGCGTCAGATGTGTATAAGAGACAG-3′) or Tn5ME-B (5′-GTCTCGTGGGCTCGGAGATGTGTATAAGAGACAG-3′) mixed to Tn5Merev (5′-[phos]CTGTCTCTTATACACATCT-3′) linker oligonucleotides, then amplified and indexed using KAPA2G Robust (Sigma). The PCR products were pooled, size selected (450-700 bp), and purified to obtain a C=[100ng/uL] and V= 30uL Novaseq x-plus sequencing sample. The sequencing was outsourced to Macrogen Europe (Supplementary Figure S4). To identify crossover sites in the examined Col × L*er mlh1* population and Col × L*er hei10-2/+* demultiplexed paired-end forward and reverse reads were aligned to the Col-0 genome reference sequence (TAIR10) using BowTie2 (44). The resultant BAM were sorted with SAMtools v1.2 (45). The identification of SNPs was carried out with SAMtools and BCFtools (46). SNP calling was based on a comprehensive list generated from a large scale of Col × L*er* population (47). The resulting tables of SNPs were filtered to retain only those exhibiting high mapping quality (>100) and high coverage (>2.5×) in R. Libraries with fewer than 50, 000 reads associated with SNPs were excluded from the analysis. Crossover calling utilized the TIGER pipeline on the filtered files (41). Finally, crossover distribution frequencies were binned into scaled windows and cumulatively aggregated across chromosome arms. A comprehensive summary of Genotyping-by-Sequencing (GBS) results is provided in Supplementary Table S4. The raw FASTQ data can be found in the NCBI Sequence Read Archive (SRA) under the BioProject accession code PRJNA1156934.

### Ploidy analysis

Raw reads were aligned to the Col-0 genome reference sequence with BowTie2 (48). The resulting BAM files were sorted and indexed with SAMtools v1.2 (49). Mosdepth was used to calculate sequencing depth with the use of -n --fast-mode -b 100000 parameters (50). Coverage was plotted for each sample and based on visual inspection; ploidy was determined.

### Cis-DCO distances

Cis-double crossover (cis-DCO) distances – defined as the observed distances between parental–heterozygous–parental genotype transitions (i.e., Col/Col– Col/L*er*–Col/Col and L*er*/L*er*–Col/L*er*–L*er*/L*er*) – were computed from GBS data. The frequency of observed distances was compared to the expected frequency of inter-crossover distances under a random crossover distribution. To obtain the expected values, 400 crossover midpoints were randomly selected from all identified crossovers within each genotype, separately for each chromosome. This process was repeated to create a second set of midpoints. For each crossover midpoint in the first set, a midpoint was randomly chosen from the second set, and the distance between them was calculated. Finally, the frequency of events, along with the median of the observed and expected cis-DCO distances, was calculated and plotted in 3.5 Mb bins.

### Quantitative RT-PCR analysis

RNA was extracted from Arabidopsis unopened flower buds (younger than stage 12) using RNeasy Mini Kit (Qiagen). cDNA was obtained using Hiscript III 1^st^ Strand cDNA Synthesis Kit + gDNA wiper (Vazyme). The expression levels of genes of interest were measured by qPCR using SYBR™ Green PCR Master Mix (Thermo) and the primers listed in Supplementary Table S5. The meiosis-specific genes *HEI10* and *DMC1* were used as a control, and *Kup9, Actin2,* and *Ubiquitin10* were used as reference.

## RESULTS

### The loss of the MutLγ complex has a less severe impact on fertility than the loss of ZMM proteins

The MutLγ complex is widely regarded as the primary resolvase responsible for crossover formation in the Class I crossover pathway (3, 51–53). Therefore, we decided to check to what extent the lack of genes encoding its components, *MLH1* and *MLH3*, affects plant fertility and crossover formation compared to *zmm* mutants. Since the available mutant alleles of *MLH1* contained either an insertion close to the 3’ end of the gene (*mlh1-1*), or in an intron (*mlh1-2*) or upstream of the alternative transcription start site (*mlh1-3*), there was a risk that these were not *null* mutants. Therefore, we used the CRISPR-Cas9 approach to generate an *mlh1-4* mutant *de novo* (Supplementary Figure S1 and S2). We measured the seed set, silique length, and pollen viability in single *mutLγ* mutants, *mlh1-1*, *mlh1-4*, *mlh3-1* and *mlh3-2,* and compared them to two well-characterized mutants of ZMM genes, *hei10* and *zip4*. While most of the mutants used for this experiment were generated in the Col-0 background, the *mlh1-1* allele originated from the Ws-2 background, therefore both accessions were used as wild-type controls. However, Col-0 and Ws-2 did not differ significantly in fertility, so we considered that *mlh1-1* could be compared with the remaining mutants in the Col background.

Visual inspection of siliques suggests that each of the *mlh1* and *mlh3* mutants is more fertile than *hei10* and *zip4*, although clearly different from wild-type plants (Figure 1A). Statistical analysis confirmed this observation (Figure 1B-D): For *mlh1* and *mlh3* mutants, from 17.4 (*mlh1-4*) to 20.7 (*mlh3-1*) seeds per silique were observed on average, and the differences between individual alleles were not statistically significant. In contrast, *zip4* and *hei10* showed only 2.3 and 5.1 seeds per silique, respectively (Figure 2B). Very similar results were also obtained for silique length (Figure 1C) and pollen viability (Figure 1D), although in these assays the *mlh1-4* allele showed a more severe fertility reduction compared to other *mlh* mutants. Interestingly, while the pollen viability in the *mlh* and *zmm* mutants was 59-73% and 24-33% of the wild type, respectively, the seed set was only 29-35% (*mlh*) and 4-8% (*zmm*) of the wild type (Figure 1D). This result is likely due to the fact that seed formation requires the simultaneous production of both functional male and female gametes. In summary, fertility comparisons show that *mutLγ* mutants have significantly higher fertility than mutants of genes encoding ZMM proteins.

**Figure 1.**
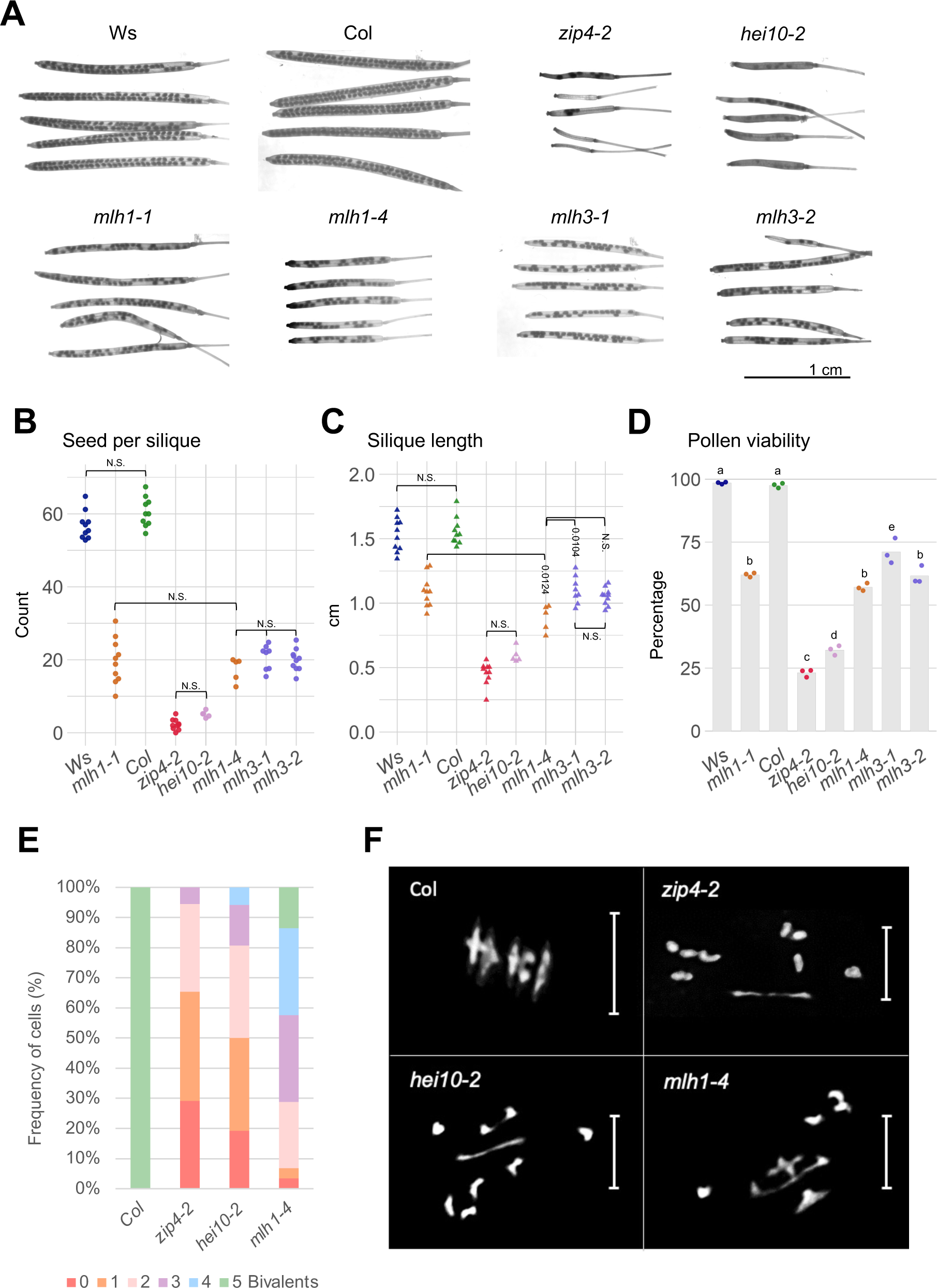
*mutLγ* mutants exhibit milder fertility and meiotic phenotypes than *zmm* mutants. **A.** Representative pictures of siliques for wild-type accessions (Ws and Col), *zmm* mutants (*zip4-2* and *hei10-2*), and *mutLγ* mutants (*mlh1-1, mlh1-4, mlh3-1,* and *mlh3-2*). Scale bar, 1cm. **B-D.** Fertility assays for *mutLγ* mutants compared to the wild types and *zmm* mutants as assessed by seed set (**B**), silique length (**C**), and pollen viability (**D**). The *P* values were estimated using one-way ANOVA and Tukey HSD tests (Supplementary Tables S10-S12). N = 5 to 10 for B-C, and n=3 for D. **E-F.** Cytological characterization of metaphase I meiocytes showing the frequency of cells with 0 to 5 bivalents (**E**), along with representative chromosome spreads from metaphase I meiocytes for Col (n = 27), *zip4-2* (n = 53)*, hei10-2* (n = 50), and *mlh1-4* (n = 57) (**F**). Scale bar, 10 μm.

**Figure 2.**
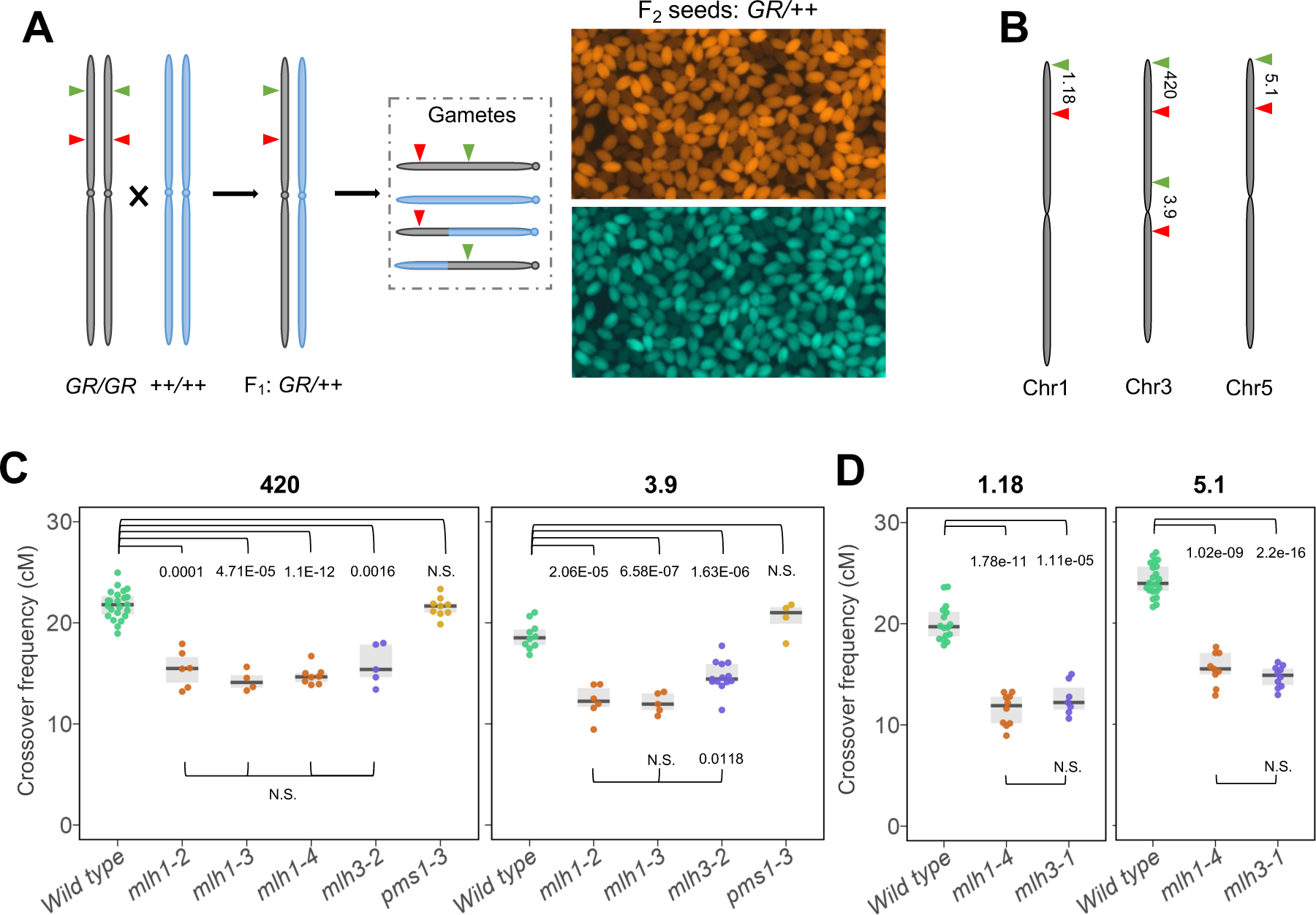
Local meiotic crossover recombination is decreased in *mutLγ* mutants. **A.** Schematic representation of crossover frequency scoring via fluorescent seed-based system. Red and green arrowheads mark the fluorescent markers delimitating a specific genomic interval. These markers are introduced into the relevant lines via crossing, and recombination frequency is determined by the segregation of the fluorescent markers. **B.** Chromosome map displaying the fluorescent intervals used in panels C-D. **C-D.** Crossover frequency (cM) for three *mlh1* alleles, two *mlh3* alleles, and one *pms1* allele in the chromosome 3 subtelomeric region *420* and pericentromeric region *3.9* (**C**), and for *mlh1-4* and *mlh3-1* in the subtelomeric regions of chromosome 1 (*1.18*), and chromosome 5 (*5.1*) (**D**). The *P* values were estimated using a Welch t-test.

Given that the dramatic decline in fertility among *zmm* mutants is attributed to a deficiency in crossover, leading to random chromosome segregation during meiosis, we sought to investigate whether crossover frequency is higher in *mlh1-4*. To this end, we examined the number of bivalents in metaphase I (Figure 1E and F, Supplementary Figure S3 and Table S6). On average, 3.19 bivalents (n = 57) were observed in the *mlh1-4* mutant, a statistically significant increase compared to 1.09 in *zip4* (n = 50; *P* = 2.2E-16, Welch t-test) and 1.70 in *hei10* (n = 53; *P* = 3.76E-12, Welch t-test). Based on these results, we concluded that *mlh1-4* mutants generate more crossovers compared to *zmm* mutants, suggesting that some Class I crossovers are independent of MutLγ.

### Local crossover frequencies are significantly reduced in *mutLγ* mutants compared to wild type

To further compare the extent of recombination reduction in *mlh* mutants, we used tester lines where the segregation of linked fluorescent reporters in seeds allows precise measurement of recombination within chromosomal intervals defined by these reporters (Figure 2A) (29, 32). For this purpose, *mlh1*, *mlh3*, *zip4*, and *hei10* mutants were crossed with the Col-*420* and Col-*3.9* lines, enabling the measurement of crossover frequencies in the subtelomeric and pericentromeric regions of chromosome 3, respectively (Figure 2B). Additionally, the *pms1* mutant was included in the analysis, as PMS1 forms the MutLα heterodimer with MLH1, and we sought to assess its role in meiotic recombination.

Measuring meiotic recombination using the fluorescent seed system was not feasible for *zip4* and *hei10* mutants due to their extremely low fertility and segregation bias. In contrast, the crossover frequency was significantly reduced in *mlh1* and *mlh3* mutants compared to the wild-type control (Figure 2C). In the *420* interval, all tested *mlh1* and *mlh3* alleles exhibited 14.3-15.9 cM, which was significantly lower than 21.8 cM for the wild type (*P* < 1.6E-03, Welch t-test). In the *3.9* interval, *mlh1* mutants showed 12.2 cM, while *mlh3* mutants showed 14.7 cM, both significantly lower than the wild type’s 18.7 cM (Figure 2C; *P* < 2.1E-05, Welch t-test). Notably, *pms1* showed no changes in crossover frequency at either interval, supporting the hypothesis that the MutLα complex does not influence meiotic recombination (Figure 2C).

To confirm that the effects of *mutLγ* mutants on crossover formation are not specific to chromosome 3 alone, we also measured recombination in *mlh1-4* and *mlh3-1* mutants at intervals *1.18* and *5.1* located on chromosomes 1 and 5, respectively (Figure 2B). In both cases, a significant decrease in the crossover rate compared to the wild type was observed, consistent with the observations on chromosome 3 (Figure 2D). In summary, *mlh1* and *mlh3* mutants exhibit a significantly reduced frequency of meiotic recombination regardless of chromosomal position, while the *pms1* mutation has no effect on crossover formation.

### The progeny of Col/L*er mlh1* mutants exhibit a notable incidence of trisomy

We aimed to assess the impact of MutLγ loss on crossover distribution across chromosomes and compare it with the distribution observed in *zmm* mutants. To achieve this, we sought to create a comprehensive genome-wide crossover map for the *mlh1* background. Since crossover sites are identified based on genotype switches, generating such a map necessitated sequencing F_2_ populations resulting from crosses between two distinct Arabidopsis accessions that are polymorphic with respect to each other. Given the absence of an *mlh1* allele in the L*er* background, we employed the CRISPR-Cas9 method to generate the *mlh1-5* mutant (see Supplementary Figure S2). Subsequently, we crossed *mlh1-4* (Col) with the *mlh1-5* (L*er*) allele and assessed fertility in the resulting F_1_ progeny. Of all the ZMM genes, *HEI10* has been shown to be dosage-dependent, and its mutant is haploinsufficient (7). Therefore, we decided to also use the F_1_ cross between *hei10* (Col) and wild-type L*er* as a control. As expected, seed set and silique length were significantly lower in the *mlh1* Col/L*er* F_1_ compared to the wild type, akin to observations in the *mlh1* Col/Col inbreds, constituting approximately 28.9% and 63.6% of wild-type values, respectively (Figure 3A and B). Although mildly affected, the seed set was also significantly lower in *hei10/+* Col/L*er*, 88.6% of the wild-type (*P* = 0.047, Welch t-test).

**Figure 3.**
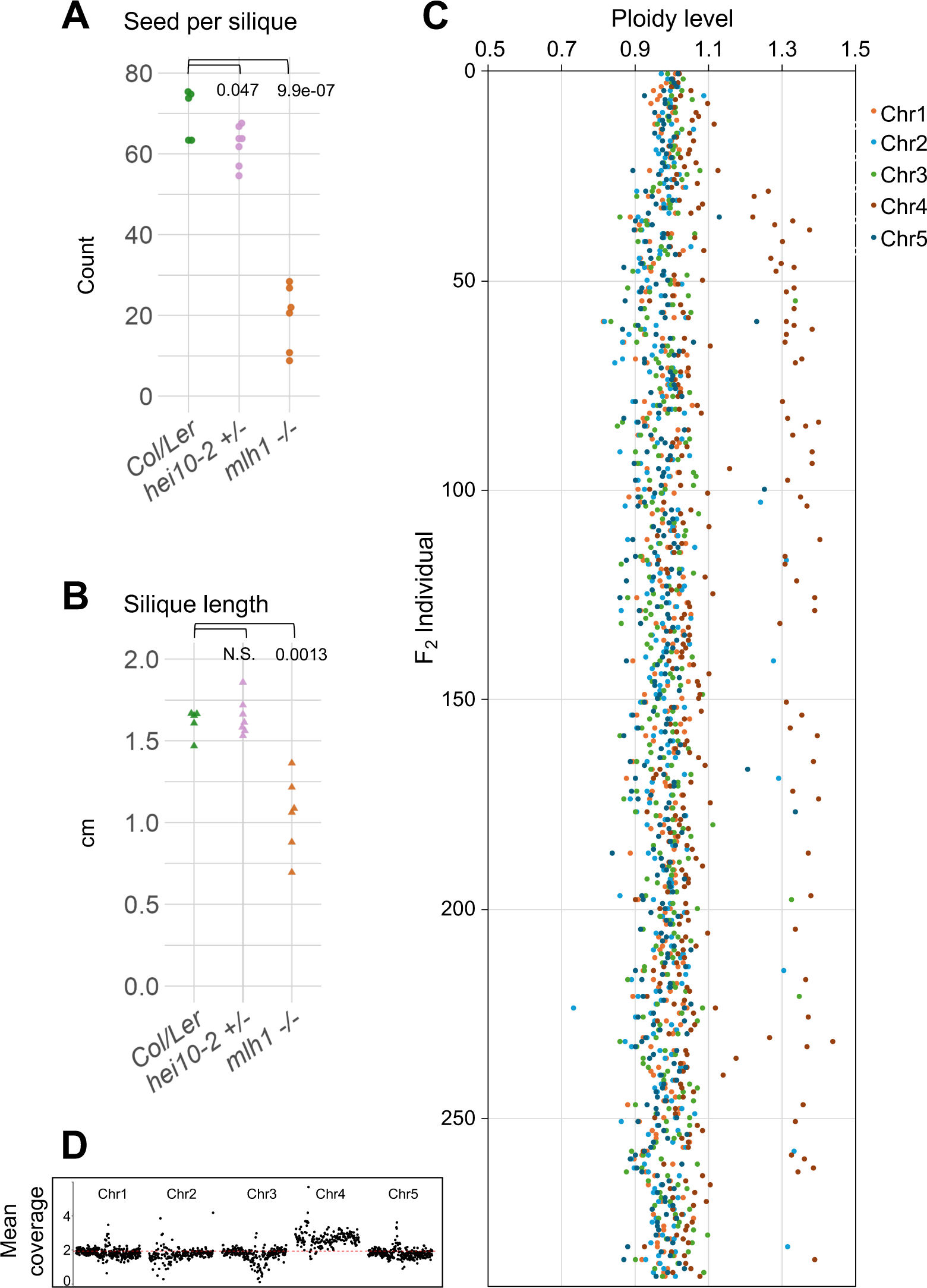
The progeny of Col/L*er mlh1* mutants exhibit a significant incidence of trisomy. **A-B.** Fertility assays for *hei10-2/+* and *mlh1* mutants compared to the wild-type Col/L*er* hybrids as assessed by seed set (**A**) and silique length (**B**). The *P* values were estimated using a Welch t-test with n = 6 to 7. **C.** Ploidy analysis for the *mlh1* Col/L*er* F_2_ population as assessed by the ploidy level for each sequenced chromosome in F_2_ individuals. **D**. Example plot illustrating trisomy of chromosome 4, detected based on mean sequencing coverage calculated in 100 kb windows.

Following self-pollination of the *mlh1* Col/L*er* F_1_ plants, we sequenced the genomic DNA of 288 descendants from a few randomly selected F_1_ individuals (Supplementary Figure S4). Similarly, we sequenced 225 F_2_ from the *hei10/+* Col/L*er* cross. Because we expected that crossover deficiency in meiosis of F_1_ plants would result in random segregation of chromosomes into gametes, we determined the level of aneuploidy in progeny. For this purpose, we examined the sequencing coverage for individual chromosomes in the F_2_ individuals and, after normalizing to the total number of reads, we calculated their ploidy level (54). We adopted a stringent criterion: a ploidy change above 1.2× indicates trisomy, while a change below 0.8× indicates monosomy for a given chromosome. We did not detect any cases of aneuploidy in *hei10/+*. In striking contrast, among 288 *mlh1* Col/L*er* F_2_ individuals, 56 plants exhibited trisomy of chromosome 4, seven plants showed trisomy of chromosome 2, three plants had trisomy of chromosome 3, and another three displayed trisomy of chromosome 5 (Figure 3C and 3D, and Supplementary Figure S5). Additionally, we observed two cases of simultaneous trisomy of chromosomes 4 and 5, one case of trisomy of chromosome 4 combined with trisomy of the left arm of chromosome 5, and four cases of partial trisomy of chromosome 4 or 5 (trisomy of an entire arm). Only one case of monosomy was found, involving chromosome 2. Interestingly, no instances of trisomy were observed for chromosome 1.

The high overrepresentation of chromosome 4 in *mlh1* aneuploids is likely due to it being the shortest chromosome, consistent with a recent report on *zyp1* and *zyp1 HEI10* lines, where trisomy was observed exclusively for chromosome 4 (9). Given the frequent occurrence of chromosome 4 trisomy compared to other types of aneuploidy in the sequenced F_2_ individuals, it is also possible that one of the F_1_ plants used was already trisomic. However, the occurrence of various types of aneuploidy is expected as a consequence of the low crossover frequency in *mlh1* mutants. An analogous analysis for the *hei10/+* mutant, which also shows a reduced crossover rate (see below), did not reveal a single case of aneuploidy. This suggests that in *mlh1*, unlike in *hei10/+*, crossover assurance is affected. We therefore conclude that the reduced recombination frequency in the Col/L*er mlh1* hybrid, in combination with reduced crossover assurance, results in a higher incidence of aneuploidy.

### Comparison of crossover distribution in *mlh1* and *hei10/+* mutants

Genotype distribution analysis along individual F_2_ chromosomes allowed us to identify the number and locations of crossovers. To avoid potential biases in crossover distribution caused by aneuploidy, we only included individuals with no evidence of ploidy changes (Figure 3C). Since only balanced gametes – primarily those that underwent crossovers on all chromosomes during meiosis – can form functional zygotes, the average crossover count estimated from F_2_ sequencing may be inflated. Despite this, the total number of crossovers across all chromosomes was significantly lower than in the wild type (Kruskal Wallis Test *P* = 0; Figure 4A). Compared to the wild type, *mlh1* mutants showed a more uniform crossover distribution along the chromosomes, with a marked reduction in recombination particularly noticeable in the pericentromeric regions, which are known for relatively high recombination activity in *A. thaliana* (Figure 4B and 4C).

**Figure 4.**
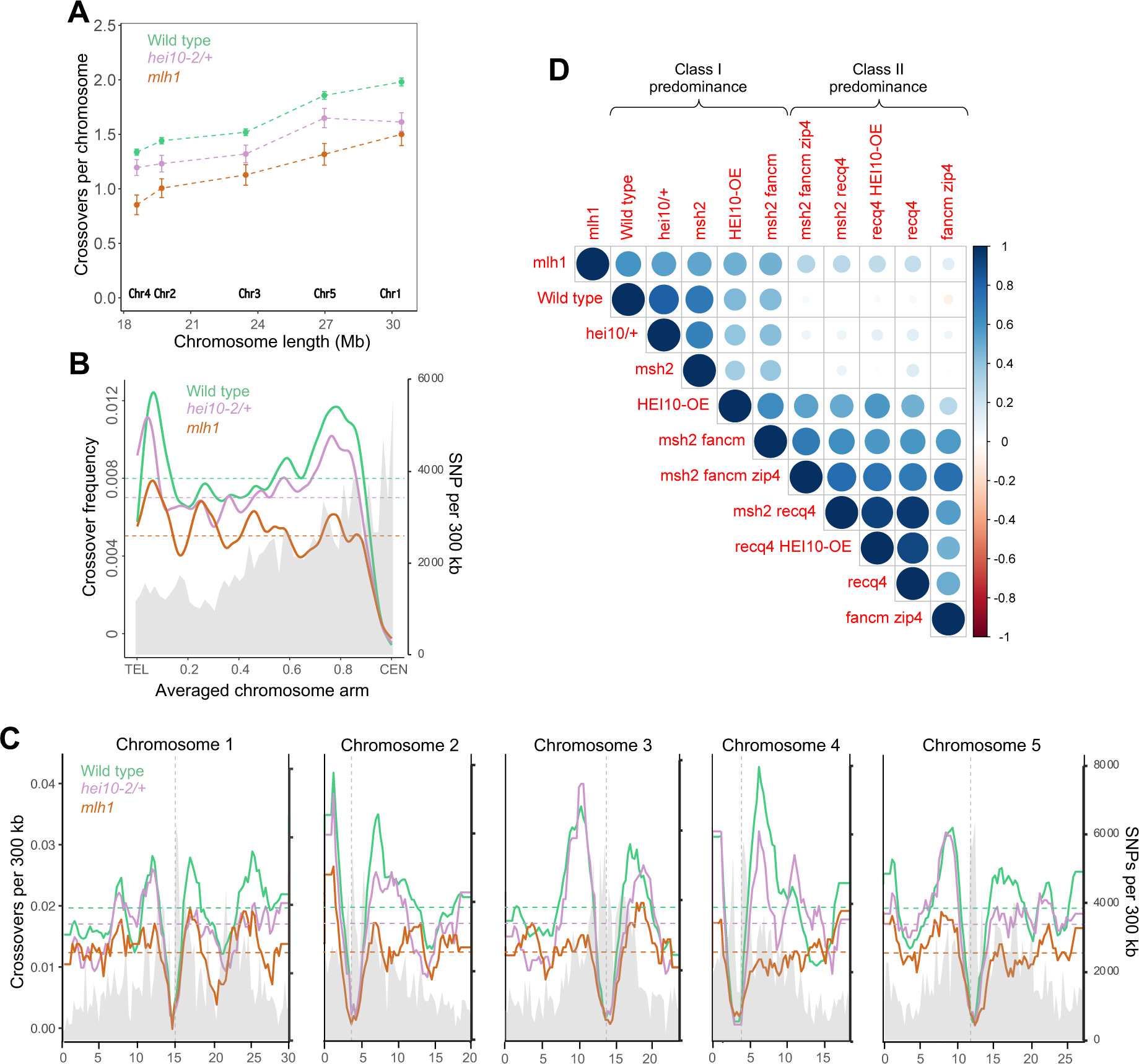
*mlh1* mutants show a reduced number of crossover events genome-wide. **A.** Differences in total crossover numbers for each chromosome in *mlh1* and *hei10/+* versus wild type. Mean crossover numbers are shown, while whiskers define SEs. **B.** Comparative representation of crossover frequency per F_2_ individual within 300 kb windows, averaged along proportionally scaled chromosome arm, orientated from telomere (TEL) to centromere (CEN). SNP density per 300 kb is shaded in grey. The mean values are shown by horizontal dashed lines. **C.** Similar to (B) but showing crossover frequency plotted along all five Arabidopsis chromosomes, averaged in 300 kb windows. Vertical dashed lines indicate centromere positions. **D.** Genome-wide correlation coefficient matrices of crossover distributions, calculated in adjacent 300 kb windows. Data for wild type, *msh2*, *HEI10-OE*, *msh2 fancm zip4*, *msh2 recq4*, *recq4 HEI10-OE*, *recq4*, and *fancm zip4* from refs. (7, 43, 47, 55, 56).

The ability to construct a crossover map for plants lacking ZMM is constrained due to the near sterility observed in *zmm* mutants (see *zip4* or *hei10* in Figure 1A and B). To gain insight into crossover distribution in ZMM-deficient plants, once again we leveraged the dosage-dependency of the *HEI10* gene. We selfed the *hei10/+* Col/L*er* plants, sequenced genomic DNA from 225 F_2_ individuals and utilized this data to construct a genome-wide crossover map. The crossover frequency was significantly reduced in *hei10/+* compared to wild-type plants (Kruskal Wallis Test *P* = 1.27E-08), although it was higher than in *mlh1* (*P* = 2.09E-06) (Figure 4A). However, the distribution of crossovers in *hei10/+* more closely resembled that in wild type (Spearman *Rho* = 0.817) than in *mlh1* (Spearman *Rho* = 0.547).

Additionally, we compared the chromosomal distribution of crossovers in *mlh1* (and *hei10/+*) with data from other genotypes (Figure 4D) (7, 43, 47, 55, 56). Correlations were significantly higher for all genotypes in which Class I crossovers predominated (*Rho* > 0.472) than for genotypes with a predominance of Class II crossovers (*Rho* ≤ 0.290). This result confirms that, despite the absence of MutLγ, the crossovers formed in the *mlh1* mutant exhibit characteristics of Class I.

### Interference is maintained but weakened in *mlh1*, increased in *hei10/+*, and negative in *msh2 fancm zip4*

The formation of each F_2_ individual results from the union of a male and a female gamete, so the analysis of F_2_ provides averaged results for both male and female meioses. However, by filtering for the parental–heterozygous–parental genotype (e.g., Col/Col–Col/L*er*–Col/Col and L*er*/L*er*–Col/L*er*–L*er*/L*er*), it is possible to identify cis-double crossovers (cis-DCOs) and measure DCO distances which inform about crossover interference (35, 57, 58). When we compared cis-DCOs in *mlh1* to the wild type, we observed that their distances were significantly shorter (Wilcoxon Signed-Rank test *P* = 6.7E-03; Figure 5A). In contrast, when we compared cis-DCOs in *hei10/+*, we found them to be significantly longer than those in the wild type (*P* = 2.4E-03; Figure 5A).

**Figure 5.**
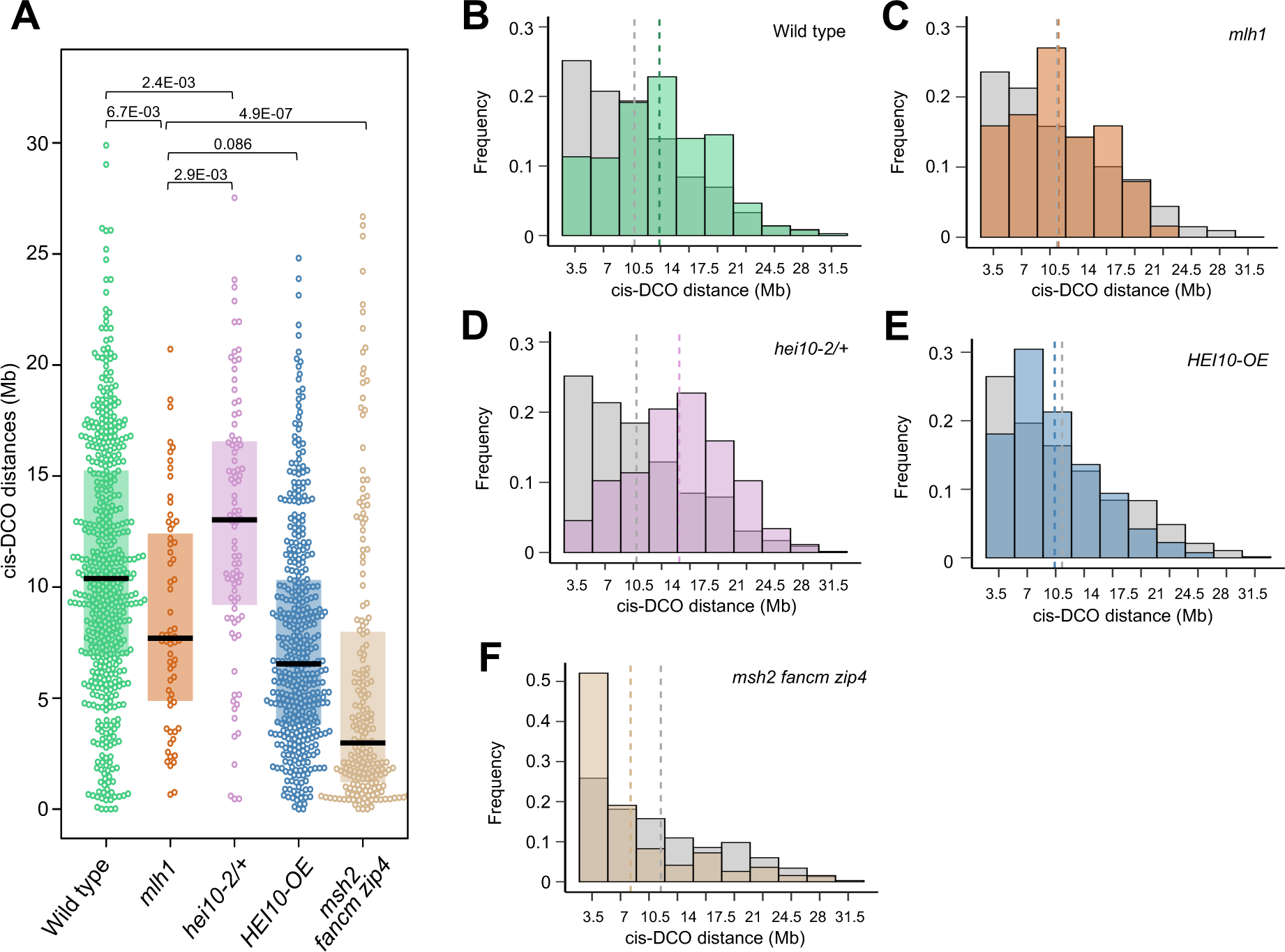
Crossover interference is decreased but maintained in *mlh1*. **A.** Cis-double crossover (cis-DCO) distances calculated from parental–heterozygous–parental genotype transitions for WT, *mlh1, hei10/+, HEI10-OE,* and *msh2 fancm zip4*. The center line of a boxplot indicates the median, while the upper and lower bounds indicate the 75th and 25th percentiles, respectively. Each dot represents one cis-intercrossover distance. The *P* values were estimated using Wilcoxon Signed-Rank test. **B-F**. Histograms illustrating the distribution of inter-crossover distances for observed data (colored bars) versus randomly generated distances (gray bars) across five genotypes: **B.** Wild type (n=1139), **C.** *mlh1* (n=63), **D.** *hei10/+* (n=88), **E.** *HEI10-OE* (n=404), and **F.** *msh2 fancm zip4* (n=194). Median inter-crossover distances for each genotype are marked with dashed lines in matching colors. Data for B, E, and F from refs. (7, 43, 47).

To examine the role of interference in the genotypes under study, we analyzed the distribution of cis-DCO distances and compared it to the expected distribution of distances between two randomly selected crossover sites. In the wild type, cis-DCOs with distances up to 7 Mb are underrepresented, while those separated by more than 10.5 Mb are overrepresented (Figure 5B). In the *mlh1* mutant, underrepresentation is also observed for cis-DCOs with distances up to 7 Mb, but overrepresentation is observed for distances greater than 7 Mb (Figure 5C). By contrary, the distribution of cis-DCO distances in *hei10/+* mirrors the wild type but is shifted further to the right, with the highest overrepresentation of DCOs separated by 14 to 17.5 Mb (Figure 5D). These findings demonstrate that although both *mlh1* and *hei10/+* mutants exhibit a reduction in crossover numbers compared to the wild type, they behave differently. In *hei10/+*, the deficiency of the pro-crossover HEI10 protein leads to increased interference, whereas in the *mlh1* mutant, interference is significantly reduced.

We also compared cis-DCO distances between the *mlh1* mutant and the *HEI10* overexpressor (*HEI10-OE*), which shows a doubled number of crossovers compared to the wild type and significantly reduced crossover interference (7, 8, 55). Interestingly, despite the stark difference in crossover numbers between *mlh1* and *HEI10-OE*, there is no significant difference in cis-DCO distances (*P* = 0.0858; Figure 5A). However, the distribution analysis of cis-DCO distances reveals that in the *HEI10* overexpression line, distances above 3.5 Mb are strongly overrepresented, which is not observed in *mlh1* (Figure 5D and E). This suggests that interference is weaker in *HEI10-OE* than in *mlh1*.

Additionally, we compared cis-DCO distances between the *mlh1* mutant and the *msh2 fancm zip4* mutant, for which we recently generated a genome-wide crossover map, although crossover interference had not yet been investigated (43). Both mutants exhibited very similar seed sets (20.08 in *msh2 fancm zip4* Col/L*er* versus 19.57 in *mlh1* Col/L*er*) and chiasma counts (3.88 in *msh2 fancm zip4* Col/Col versus 3.75 in *mlh1* Col/Col). However, in *msh2 fancm zip4*, Class I crossovers are entirely absent due to the loss of the key ZMM protein ZIP4 (43, 59). The cis-DCO distances were significantly shorter in *msh2 fancm zip4* than in *mlh1* (median = 2.98 Mb versus 7.70 Mb, Wilcoxon Signed-Rank test *P* = 4.9E-07; Figure 5A). Unexpectedly, the frequency distribution of cis-DCO distances in *msh2 fancm zip4* revealed a pronounced overrepresentation of the shortest distances below 3.15 Mb, occurring at twice the expected frequency for a random distribution (Figure 5F). This indicates that crossover interference in the *msh2 fancm zip4* triple mutant is negative, meaning that crossovers tend to cluster together. The likely cause of this phenomenon lies in the severely altered crossover distribution, which is concentrated almost exclusively at the chromosome ends. Combined with the absence of Class I crossovers, this dramatically increases the likelihood of closely spaced events beyond what would be expected in a random distribution.

In summary, our results suggest that MutLγ complexes are not essential for maintaining interference, and the reduced interference observed in *mlh1* is likely due to a proportionally higher number of Class II crossovers compared to the wild type. Similarly, interference decreases in the line overexpressing the pro-crossover factor HEI10, but this reduction is attributed to an increased number of crossovers. Conversely, in *HEI10* haploinsufficient lines (*hei10/+*), interference increases due to a reduced number of crossovers; however, there are still enough Class I events present that Class II crossovers do not significantly affect the overall distribution of recombination, thus enhancing interference. In the *msh2 fancm zip4* mutant, only Class II crossovers occur, which tend to cluster due to a distribution of events that is highly skewed toward the chromosome ends.

### Concomitant loss of MutLγ and MUS81 leads to a significant decrease in fertility, chiasma frequency and crossover assurance

Due to the significantly higher chiasma number and fertility of *mutLγ* mutants compared to *zmm* mutants, and the distinct recombination distribution patterns in *mlh1* versus *hei10/+* plants, it appears that another endonuclease partially compensates for the loss of MutLγ. To investigate this, we generated the *mlh1 mlh3 mus81* triple mutant and compared its fertility with other mutants (Figure 6A and B, and Supplementary Figure S6A). Seed set analysis revealed that *mlh1 mlh3 mus81* produces significantly fewer seeds than the *mlh1 mlh3* double mutant (mean = 4.69 and 18.00, respectively; *P* = 1.45E-02, one-way ANOVA and Tukey HSD). Notably, we also observed a slight reduction in seed set in *mus81* compared to the wild-type Col (mean = 51.4 and 60.0, respectively; *P* = 0.021; Figure 6A) (22). Similar trends were observed in the analysis of silique length across *mus81*, *mlh1*, *mlh3* mutants, and their combinations (Supplementary Figure S6A). These findings demonstrate that MUS81 plays a critical role when the absence of MutLγ disrupts the proper functioning of the ZMM pathway.

**Figure 6.**
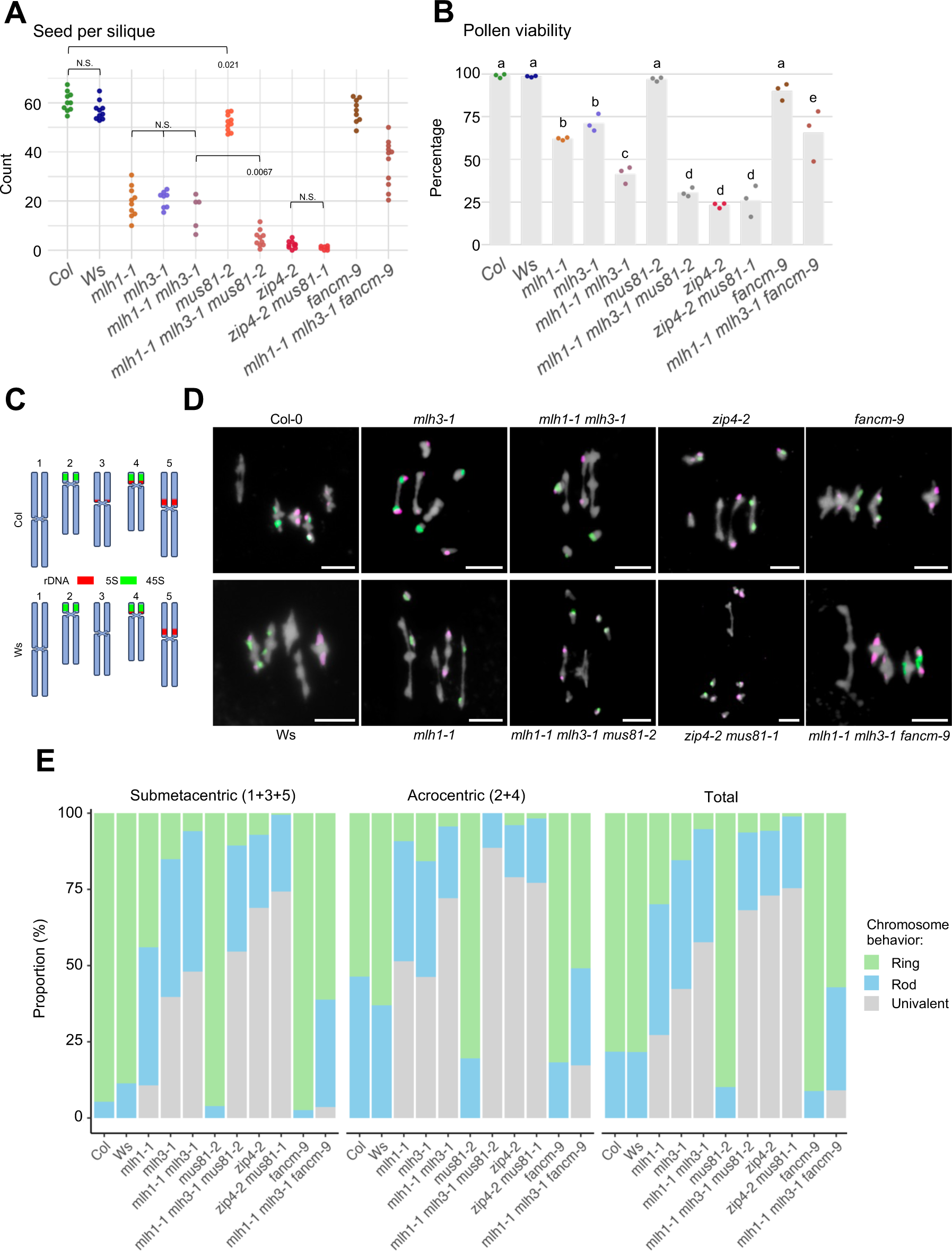
MUS81 endonuclease partially compensates for the loss of MutLγ. **A-B.** Fertility as assessed by seed set (**A**) and pollen viability (**B**). The *P* values were estimated using one-way ANOVA and the Tukey HSD tests (Supplementary Tables S13-S14). Sample sizes: n = 5 to 11 in A, and n = 3 in B. **C-E.** Cytological characterization of metaphase I meiocytes. **C**. Chromosome representations of 5S and 45S rDNA positions in Col and Ws genetic backgrounds. **D.** Representative pictures of chromosome spreads of metaphase I meiocytes stained with DAPI and labeled with FISH against 5S (red) and 45S (green) rDNA. Scale bar, 10 μm. **E.** Frequency of ring, rod, and univalent chromosome configuration in submetacentric chromosomes (1, 3 and 5), acrocentric chromosomes (2 and 4), and the average across all chromosomes. The number of characterized meiocytes for each genotype is as follows: Col (n = 69), Ws (n = 50)*, mlh1-1* (n = 53)*, mlh3-1* (n = 108), *mlh1-1 mlh3-1* (n = 34), *mus81-2* (n = 102), *mlh1-1 mlh3-1 mus81-2* (n = 44), *zip4-2* (n = 255), *zip4-2 mus81-1* (n=57), *fancm-9* (n = 52), and *mlh1-1 mlh3-1 fancm-9* (n = 55).

On the other hand, *mlh1 mlh3 mus81* did not produce significantly more seeds than *zip4* (mean = 2.27) and *zip4 mus81* (mean = 0.93) (Figure 6A). However, drawing conclusions from the comparison of these genotypes is subject to inaccuracy due to very low seed number values. Similarly, silique length shows statistically insignificant differences between these genotypes. Because seeds are produced by the fusion of two gametes, low seed set and silique length values are the product of reduced viability of both the male and female gametes. Therefore, we also examined pollen viability, the decrease of which in meiotic mutants is a consequence exclusively of impaired male meiosis. We did not observe statistically significant differences between the *mlh1 mlh3 mus81*, *zip4*, and *zip4 mus81* mutants (Figure 6B). The lack of expected differences between *zip4* and *zip4 mus81* indicates that this analysis is also not precise enough to determine whether MUS81 can contribute to ZMM-mediated crossover repair when MutLγ is absent.

To confirm that the observed decrease in fertility in the *mlh1 mlh3 mus81* mutant compared to *mlh1 mlh3* is due to a further reduction in the number of crossovers, we analyzed bivalent configurations at metaphase I. We used fluorescence in situ hybridization (FISH) with 5S and 45S rDNA probes to distinguish submetacentric chromosomes (1, 3, and 5) from acrocentric chromosomes (2 and 4) (Figure 6C and D). This approach enabled us to assess how often individual chromosomes fail to form a chiasma. We evaluated bivalent formation and chiasma count in single, double, and triple mutants (Figure 6D and E, and Supplementary Figure S6B and C, and Supplementary Tables S7, S8, and S9). In the *mlh1 mlh3 mus81* triple mutant, we observed a higher total number of univalents than in the *mlh1 mlh3* mutant, which itself had a greater frequency of univalents than the single mutants (Figure 6E). Notably, the frequency of univalents on acrocentric chromosomes increased by 16.5%, from 72.1% in *mlh1 mlh3* to 88.6% in *mlh1 mlh3 mus81*. In contrast, the increase in univalents on submetacentric chromosomes was only 6.5%, from 48.0% to 54.5% (Figure 6E and Supplementary Tables S8 and S9). Additionally, no ring bivalents were observed on acrocentric chromosomes in the triple mutant. By comparison, *mus81* and *pms1* mutants showed no significant differences from the control in terms of bivalent configurations and chiasma counting (Figure 6E and Supplementary Figure S6C). This suggests that in the *mlh1 mlh3 mus81* background, crossover assurance is weakened compared to *mlh1 mlh3*. Since crossover assurance is a hallmark of Class I (ZMM-dependent) crossovers, this result suggests that MUS81 may partially take on the role of resolvase within the ZMM pathway when MutLγ is not available.

The *mlh1 mlh3 mus81* mutant exhibited fewer univalents than both *zip4* and *mus81 zip4* (Figure 6E). This suggests that MUS81 is not the only endonuclease capable of resolving ZMM-dependent recombination intermediates into crossovers when MutLγ proteins are absent. Interestingly, this difference in univalent frequency was observed in submetacentric chromosomes, while for acrocentric chromosomes, we observed the opposite effect: the triple mutant had more univalents than *zip4* and *mus81 zip4* (Figure 6E and Supplementary Tables S8 and S9). These results suggests that crossover assurance is more compromised in *mlh1 mlh3 mus81* compared to *zip4* and *mus81 zip4*.

### Knocking out *FANCM* partially improves bivalent formation and rescues the low fertility phenotype in *mutLγ* mutants

The low fertility of *zmm* mutants is caused by a deficiency in crossovers due to the complete disruption of the ZMM crossover pathway. Increasing the frequency of ZMM-independent Class II crossovers can restore fertility. In wild-type Arabidopsis, Class II crossovers are rarely formed because their substrates are typically removed by the activity of DNA helicases like FANCM and RECQ4. Inactivation of these helicases leads to a significant increase in Class II crossovers. Consequently, in *zmm* mutants, the simultaneous loss of FANCM or RECQ4 substantially improves fertility (27, 28, 60).

To test whether the fertility of MutLγ complex mutants could be similarly improved, we generated the *mlh1 mlh3 fancm* triple mutant and compared it with both the *fancm* single mutant and the *mlh1 mlh3* double mutant. The triple mutant produced an average of 35.8 seeds per silique, significantly more than the 15.7 seeds of the *mlh1 mlh3* mutant (*P* = 6.98E-8, one-way ANOVA and Tukey HSD; Figure 6A), although still fewer than the wild type (60.5 seeds per silique; *P* = 1.43E-11). A similar increase in fertility has been observed in the *zip4 fancm* mutant (28). We also examined silique length and pollen viability; these results were consistent with the conclusion that the mutation in *fancm* rescues the fertility of *mutLγ* mutants, as it does for *zmm* mutants (Figure 6B and Supplementary Figure S6A).

We then identified individual chromosomes using FISH and examined the bivalent configurations at metaphase I. The *mlh1 mlh3 fancm* triple mutant exhibited over six times fewer univalents and more than ten times as many ring bivalents compared to the *mlh1 mlh3* double mutant, indicating a significant increase in crossover frequency (Figure 6E and Supplementary Figure S6C and Tables S7, S8 and S9). Interestingly, univalents were over four times more common on acrocentric chromosomes compared to submetacentric ones in *the mlh1 mlh3 fancm* mutant, accounting for 17.3% and 3.6%, respectively (Figure 6E and Supplementary Tables S8 and S9). This suggests a lack of crossover assurance, consistent with the observed increase in Class II crossovers in this mutant.

Overall, these data show that, similar to *zmm* mutants, the loss of FANCM in *mutLγ* mutants also increases crossover frequency, which contributes to their improved fertility. However, despite this increase, the triple mutant had a lower frequency of ring bivalents (57.1%) compared to the control (78.3%), and univalents persisted. This suggests that, unlike in *zmm* mutants, the *fancm* mutation does not fully restore the chromosomal configurations seen in the control in the *mlh1 mlh3* background.

### Elevated expression of *MLH1* or *MLH3* causes a moderate increase in crossover frequency

We sought to determine whether increasing the expression of *MLH1*, *MLH3*, and *PMS1* would affect crossover frequency. To this end, we cloned the genomic sequences of these genes under their native promoters and transformed them individually into Col-*420* plants hemizygous for fluorescent crossover reporters (Supplementary Figure S7). T_1_ plants segregating for the *420* reporters were used to assess crossover frequency within this interval. Most T_1_ plants exhibited crossover frequencies similar to the wild type, with no significant changes observed in the T_2_ generation following selfing (Supplementary Figure S9A and B). For selected *pMLH1::MLH1* and *pMLH3::MLH3* T_2_ plants, we isolated RNA from closed flower buds and measured gene expression levels using RT-qPCR. We then correlated these levels with the *420* crossover frequency. In *pMLH3::MLH3* T_2_ plants, a positive correlation was observed (Spearman *Rho* = 0.714; Supplementary Figure S9C). However, we also noted that some T_2_ plants with increased expression exhibited reduced fertility (Supplementary Figure S9D and E).

In parallel, three independent T_2_ plants for both MLH1 and MLH3 constructs were crossed with Col-*420* or L*er* to generate F_1_ inbred (Col × Col*-420*) and F_1_ hybrid (Col*-420* × L*er*) progeny (Supplementary Figure S7). Crossover frequency measurements in the *420* interval revealed that lines carrying additional copies of *MLH1* exhibited a significant increase in recombination in the inbred background, though this effect was not observed in hybrids with L*er* (Figure 7A). We also repeated this experiment for the *3.9* interval, which allows measurement of crossover frequency in the pericentromeric interval (the location is shown in Figure 2B), and obtained very similar results (Figure 7B). Interestingly, a significant increase in recombination frequency was observed in both inbred and hybrid backgrounds for lines with additional copies of *MLH3* (Figure 7A), suggesting that elevated *MLH3* expression enhances crossover rates regardless of genetic background.

**Figure 7.**
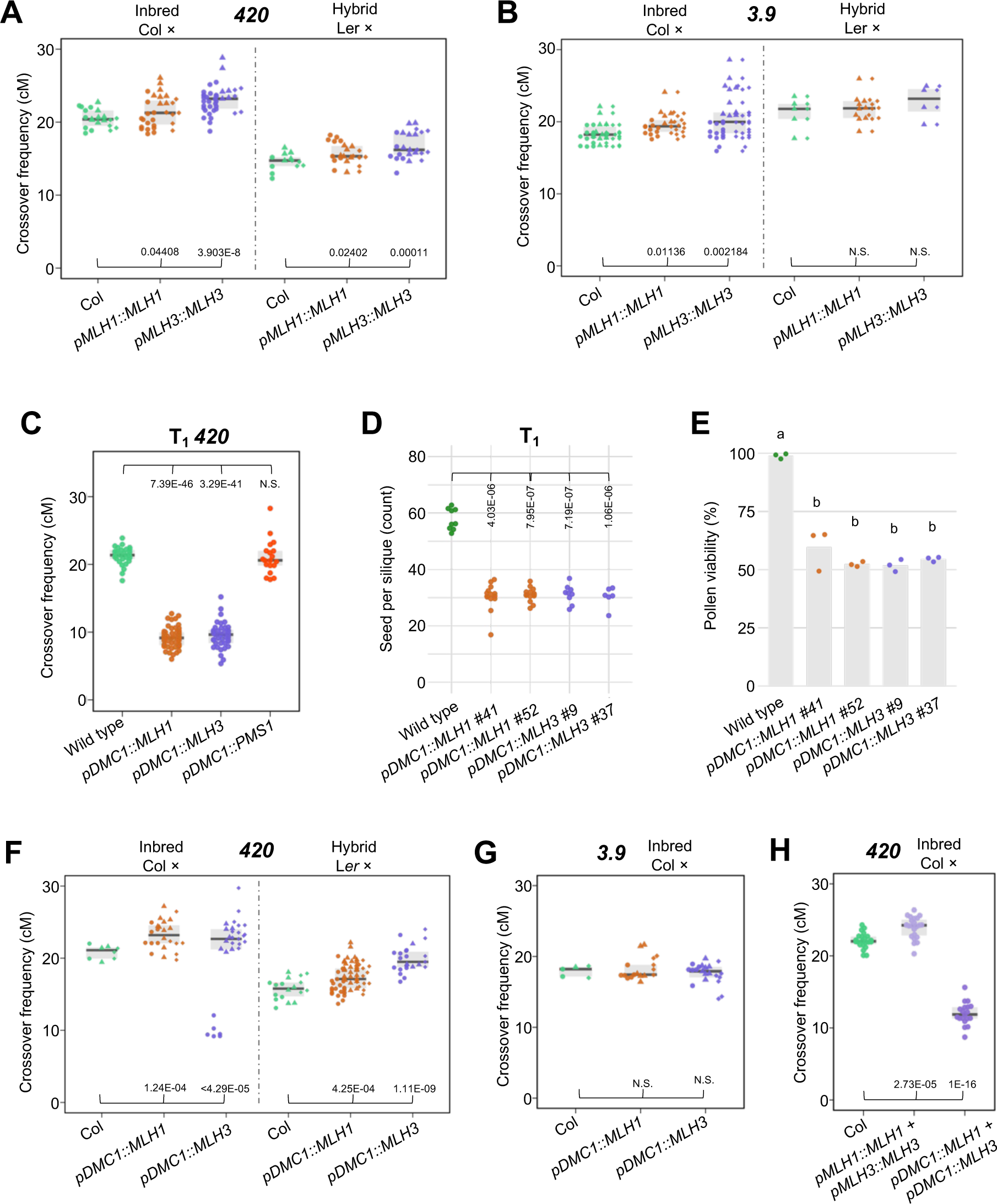
Increased MutLγ expression boosts crossovers, but overexpression drastically reduces crossover numbers and plant fertility. **A.** Crossover frequency in inbred Col/Col and hybrid Col/L*er* F_1_ plants with additional *MLH1* or *MLH3* copies under their respective native promoters, measured in *420* subtelomeric region of chromosome 3. Each datapoint represents a measurement from one plant. Independent transformant lines for each genotype are indicated by a different datapoint shape. **B.** As for (A) but measured in *3.9* pericentromeric region. **C.** Crossover frequency of *MLH1, MLH3,* and *PMS1* overexpressors under the control of *DMC1* promoter at the T_1_ generation, measured in *420*. Each datapoint represents a measurement from one plant. **D.** Fertility assays for *MLH1* and *MLH3* overexpressors, under *DMC1* promoter, assessed by seed set. Each data point represents the average of five siliques from a single plant. The *P* values were estimated using one-way ANOVA and the Tukey HSD tests (Supplementary Table S17). **E.** As in (D) but showing pollen viability. Each datapoint represents a measurement of 500 pollen grains from one plant. The *P* values were estimated using one-way ANOVA and the Tukey HSD tests (Supplementary Table S18). **F-G.** Crossover frequency in inbred Col/Col and hybrid Col/L*er* F_1_ plants overexpressing additional *MLH1* or *MLH3* copies under *DMC1* promoter, measured in the *420* interval (**F**) and in Col/Col inbred plants in the 3.9 interval (**G**). **H.** Crossover frequency in inbred Col/Col F_1_ plants with additional *MLH1* and *MLH3* copies (double overexpressors) under their native or *DMC1* promoter, measured in the *420* interval. For crossover frequency (**A-C, F-H**) the *P* values were estimated using the Welch t-test.

### Excessive overexpression of *MLH1* or *MLH3* inhibits crossover recombination and impacts fertility

Since some T_1_ plants showing increased expression of *MLH1* or *MLH3* also exhibited reduced fertility (Supplementary Figure S9D and E), we hypothesized that an excess of these proteins might be toxic for the cell. To verify this hypothesis, we modified the genetic constructs for the *MLH1*, *MLH3* and *PMS1* genes by substituting their promoters for the promoter of the *DMC1*. The DMC1 recombinase serves as a meiosis-specific RAD51 homolog, exhibiting expression levels approximately 100 times higher than Arabidopsis MLH3 (Supplementary Figure S8). Subsequently, we transformed *420* segregating plants with these modified constructs and assessed crossover frequency. Remarkably, virtually all T_1_ plants carrying the *pDMC1::MLH1* and *pDMC1::MLH3* constructs displayed a significantly reduced crossover frequency, plummeting from ∼22 cM in the wild type to only ∼10 cM in T_1_ (Figure 7C). Conversely, no such effect was observed for plants with the *pDMC1::PMS1* construct, confirming the negligible role of PMS1 in meiotic recombination (Figure 7C).

The T_2_ generation plants also showed a reduced *420* crossover frequency (Supplementary Figure S10A). Additionally, fertility assessments conducted on T_2_ plants carrying the *pDMC1::MLH1* and *pDMC1::MLH3* constructs revealed a notable decrease in seed set and pollen viability, dropping to approximately 50% of the level observed in the wild type (Figure 7D and E). We did not observe any phenotypic changes at the level of vegetative development or in the structure of flowers. This is consistent with the meiosis-specific activity of the *DMC1* promoter and suggests that the overexpression of *MLH1* and *MLH3* manifests in disturbances exclusively during the meiotic stage. These findings underscore the toxic effects of excessive expression levels of the MutLγ complex genes on plant fertility.

Three independent T_2_ plants were subsequently crossed with Col-*420* and L*er* to assess recombination in both inbreds and hybrids (Supplementary Figure S7). In the resulting F_1_ plants, the *420* crossover frequency was measured, which, in some instances, surpassed that of the wild-type control (Figure 7F and G). We argue that by crossing with a wild-type plant, the supply of the protein is reduced by half, which in many lines brings its level back to a point that does not cause a decrease in crossover frequency and the associated drop in fertility. Notably, in one line carrying the *pDMC1::MLH3* construct, recombination decreased to 10 cM in inbred, akin to the level observed in self-pollinated T_2_ plants. However, this same line exhibited an increased *420* crossover frequency in crosses with L*er* (Figure 6E; mean 19 cM versus 15 cM in the wild type, *P* = 0.0041, Welch t-test). Such evident alterations in crossover frequency underscore the sensitivity of MutLγ components to expression levels.

Because the MutLγ complex is a heterodimer containing both MLH1 and MLH3, we decided to investigate how meiotic recombination would be affected by simultaneous increase in the expression of both genes. To this end, we crossed T_2_ plants carrying additional copies of *MLH1* and *MLH3*. Due to the reduced fertility of T_2_, only a small proportion of crosses were successful. Pooled F_1_ from three crosses between independent T_2_ plants with additional copies of *MLH1* and *MLH3* under their native promoters showed an increased *420* crossover frequency (Figure 7H; 23.94 cM versus 22.11 cM for wild-type control; *P* = 1.27E-05, Welch t-test). However, in double overexpressor plants with additional genes driven by the *DMC1* promoter, recombination was significantly lower than in the wild type, as was the case in single overexpressors (Figure 7H; 11.97 cM; *P* = 6.72E-22). These results show that simultaneous upregulation of both genes of the MutLγ complex does not result in significant changes in recombination rates compared to those observed in single overexpressors. In conclusion, modifying the expression of MutL complex genes cannot be routinely used to increase recombination frequency because plants are highly sensitive to the levels of MLH1 and MLH3 in the cell. Excessive expression is often toxic and can lead to a drastic decline in fertility.

## DISCUSSION

The MutLγ complex is widely regarded as the main resolvase for Class I crossovers, which depend on ZMM proteins (hence the name ZMM crossover pathway) (3, 53, 61). Despite their role in this pathway, MLH1 and MLH3 are not classified as ZMM proteins, and their role in the pathway, aside from functioning as a resolvase, remains unclear. The primary reason for excluding MutLγ from the ZMM group lies in the distinct phenotypic differences: in many eukaryotes, mutants lacking components of the MutLγ complex exhibit a milder phenotype compared to classical ZMM mutants, displaying higher fertility and an increased crossover frequency (62–67). In this study, we undertook a comprehensive characterization of the MutLγ complex in *A. thaliana*, employing a diverse set of methodologies.

We analyzed plant fertility and bivalent formation during meiosis in various alleles of *mlh1* and *mlh3*, including the *mlh1-4* null allele generated using CRISPR-Cas9 technology (this study). As expected, all mutants showed reduced fertility and fewer bivalents than the wild type, but the defects were less severe than in the ZMM mutants *zip4* and *hei10* (Figure 1) (15, 16, 68). Fluorescent seed-based measurement of crossover frequency in chromosomal intervals showed a decrease in recombination in both pericentromeric and subtelomeric regions in *mlh1* and *mlh3* mutants (Figure 2). Notably, the *pms1* mutant, a component of the MutLα complex, did not show significant changes compared to the wild type, confirming that it does not play a significant role in meiotic recombination (Figure 2) (20).

We also generated genome-wide crossover maps for *mlh1* to assess its similarity to ZMM mutants. To further investigate this, we constructed a crossover map for the *hei10/+* mutant, the only ZMM mutant with a dosage effect (as homozygous mutants are nearly infertile, making crossover analysis unreliable) (7, 69). Since reduced crossover numbers can lead to aneuploidy, we first examined the ploidy levels in F_2_ plants through sequencing. While no aneuploids were found in *hei10/+*, a significant number were present in *mlh1* F_2_ plants (Figure 3C and D). The *mlh1* mutant exhibits a different recombination pattern compared to *hei10/+* and wild type, showing a more even distribution of crossovers along chromosomes, particularly noticeable along the chromosomal arms (Figure 4B and C). In contrast, crossover patterns in *hei10/+* are similar to wild type, with pronounced increases in pericentromeric and subtelomeric regions (Figure 4B and C). We further compared these patterns with other genotypes analyzed through the same method (Figure 4D) (7, 43, 47, 55, 70). These comparisons revealed that the crossover distribution in *mlh1* resembles Class I crossover patterns rather than Class II. This suggests that many crossovers in *mlh1* exhibit Class I characteristics, and the observed deviations from wild-type distribution may stem from a reduced Class I/Class II ratio.

We conducted a crossover interference analysis based on cis-DCO distances, which measure the spacing between two crossovers occurring on the same chromosome during meiosis. Shorter distances between crossover pairs indicate weaker interference (57). In addition to our data for *mlh1* and *hei10/+*, we analyzed previously published datasets (not yet evaluated for cis-DCO distances) for lines overexpressing *HEI10*, as well as the *msh2 fancm zip4* mutant and wild type (7, 43, 47). The *hei10/+* mutant showed increased interference compared to wild type (Figure 5A and D), consistent with the known pro-crossover role of HEI10 and the coarsening model of crossover interference. In this model, reduced HEI10 availability limits the number of HEI10 foci that can mature into Class I crossovers, thus enhancing interference (8, 10). Conversely, *HEI10* overexpression led to decreased interference, as more HEI10 loci matured into crossover-competent foci (Figure 5A and E), aligning with previously published results on crossover interference in this line (8, 55).

The *mlh1* mutant exhibited significantly shorter cis-DCO distances compared to wild type, indicating reduced interference (Figure 5). Notably, cis-DCO distances in *mlh1* were not significantly different from those in the *HEI10*-overexpressing line, despite an approximately four-fold difference in crossover numbers between these genotypes. This suggests that the mechanism of interference reduction in *mlh1* is distinct from that in *HEI10* overexpression. In *mlh1*, the reduction appears to stem from the loss of the primary resolvase in the ZMM pathway, while other pathway components remain intact (8, 55). We propose that the main cause of reduced interference in *mlh1* is the altered Class I/Class II crossover ratio, driven by the drastic reduction in Class I crossovers.

Based on this conclusion, we hypothesized that interference in the *mlh1* mutant is not abolished because some recombination intermediates, protected by ZMM proteins, are resolved into crossovers by alternative nucleases. To test this hypothesis, we compared cis-DCO distances in the *mlh1* mutant with those in the *msh2 fancm zip4* mutant. In this latter mutant, Class I crossovers cannot form due to the *zip4* mutation, while Class II crossovers are elevated by the loss of the anti-crossover helicase FANCM and inactivation of the mismatch sensor MSH2 (28, 43, 59). Interestingly, despite both mutants exhibit similar fertility and crossover numbers, the *msh2 fancm zip4* mutant showed significantly shorter cis-DCO distances than *mlh1* (Figure 5). This difference arises because Class II crossovers in *msh2 fancm zip4* are not subject to interference. These results support the idea that while crossover interference is reduced in the *mlh1* mutant, it remains functional.

We also performed extensive genetic interaction studies involving *mlh1* and *mlh3* in combination with other meiotic recombination mutants. By analyzing fertility and chiasma numbers, we confirmed that another nuclease partially compensates for the loss of MutLγ in mutants lacking MLH1 and/or MLH3 (Figure 6). Higgins et al. (4) observed that prophase I progression is delayed threefold longer in *Atmlh3* compared to *Atmsh4*, suggesting that this extended prophase might provide more time for an alternative nuclease to repair ZMM-protected recombination intermediates through crossovers. Importantly, the *mlh1 mlh3 mus81* triple mutant shows significantly weaker crossover assurance compared to the *mlh1 mlh3* double mutant, as deduced from the substantial increase in the number of univalents on acrocentric chromosomes compared to submetacentric chromosomes (Figure 6E). This suggests that MUS81 takes on the role of resolvase in the absence of MutLγ (Figure 6E). Crossover assurance remains intact in *mlh1 mlh3* mutants, as ZMM proteins continue to designate recombination intermediates for crossover formation. However, unlike MutLγ, MUS81 resolves these intermediates with equal likelihood as either crossovers or non-crossovers, which ultimately weakens crossover assurance. Moreover, the differences between *mlh1 mlh3 mus81* and *mus81 zip4* suggest that MUS81 is not the only nuclease involved in compensating for the loss of MutLγ. This result should be taken with caution because the low chiasma numbers and fertility measurements in the compared mutants are prone to error.

We also found that the mutation of the anti-crossover DNA helicase FANCM significantly mitigates the effects of MutLγ loss (Figure 6). It is already established that FANCM suppresses mutations in *zip4* and other ZMM genes (28, 60, 71–73). However, the findings regarding the mutant lacking MutLγ are particularly noteworthy because the ZMM complexes remain fully functional, with only the main resolvase absent. It is widely accepted that ZMM proteins protect recombination intermediates from the actions of DNA helicases and structure-specific nucleases (3, 74, 75). Thus, we conclude that in the presence of ZMM proteins and in the absence of MutLγ, Class II crossovers can still occur in regions occupied by ZMM.

We also examined how increasing the expression of MutLγ components influences crossover frequency. A moderate increase in *MLH3* expression resulted in a modest rise in crossover frequency in both subtelomeric and pericentromeric regions (Figure 7). The effect of MLH1 was less pronounced, indicating that MLH3 may be the limiting factor within the MutLγ complex. This is consistent with the significantly lower native expression of *MLH3* compared to *MLH1*, as well as the meiosis-specific nature of MLH3 (Supplementary Figure S8) (15). Additionally, the meiotic phenotypes of the single mutants differ, with *mlh3* exhibiting a higher frequency of univalents and rod bivalents compared to *mlh1* (Figure 6). However, excessive and premature expression of either *MLH1* or *MLH3* – unlike *PMS1* – resulted in a drastic reduction in both crossover frequency and plant fertility. Recent studies in yeast have revealed an unexpected interaction between MLH1-MLH3 and the recombinase DMC1 (76). It is possible that premature overexpression of MutLγ components leads to competition with DMC1, thereby restricting proper strand invasion and/or the formation of early recombination intermediates. This highlights the need for strict regulation of MLH1 and MLH3 protein levels during meiosis. In contrast, our findings suggest that PMS1 does not play a significant role in meiotic recombination.

## Data availability

The data supporting this article can be accessed in the published version or its supplementary files. Additionally, raw genome sequencing data for the mutant *mlh1* and *hei10/+* Col×L*er* F_2_ populations are deposited in the NCBI Sequence Read Archive (SRA) under the BioProject accession code PRJNA1156934.

## Acknowledgments

We thank Avi Levy (The Weizmann Institute, Israel) and Scott R. Poethig (University of Pennsylvania, US) for FTLs (CTLs); François Belzile (Université Laval, Canada) for sharing *mlh1-1* and *pms1-1*, Raphaël Mercier (Max Planck Institute for Plant Breeding Research, Germany) for sharing *mlh1-3*; B. Martín, M.C. Moreno, J. Barrios and H. Korcz-Szatkowska for their excellent technical assistance.

## Author contribution

NK, JLS, MP, and PAZ designed the research; NK, NFJ, WD, AP, ESZ, and AMV performed the experiments; WD performed the computational analyses, NK, NFJ, WD, AP, MP, and PAZ analyzed the data; and NK, MP, and PAZ wrote the article with aid of all authors.

## References

1. Villeneuve, A.M. and Hillers, K.J. (2001) Whence meiosis? Cell, 106, 647–650.

2. McDonald, M.J., Rice, D.P. and Desai, M.M. (2016) Sex speeds adaptation by altering the dynamics of molecular evolution. Nature, 531, 233–236.

3. Hunter, N. (2015) Meiotic recombination: The essence of heredity. Cold Spring Harb Perspect Biol, 7, a016618.

4. Higgins, J.D., Armstrong, S.J., Franklin, F.C.H. and Jones, G.H. (2004) The Arabidopsis MutS homolog AtMSH4 functions at an early step in recombination: evidence for two classes of recombination in Arabidopsis. Genes Dev, 18, 2557–2570.

5. France, M.G., Enderle, J., Röhrig, S., Puchta, H., Franklin, F.C.H. and Higgins, J.D. (2021) ZYP1 is required for obligate cross-over formation and cross-over interference in Arabidopsis. Proc Natl Acad Sci U S A, 118, 1–11.

6. Capilla-Pérez, L., Durand, S., Hurel, A., Lian, Q., Chambon, A., Taochy, C., Solier, V., Grelon, M. and Mercier, R. (2021) The synaptonemal complex imposes crossover interference and heterochiasmy in Arabidopsis. Proc Natl Acad Sci U S A, 118, 1–11.

7. Ziolkowski, P.A., Underwood, C.J., Lambing, C., Martinez-Garcia, M., Lawrence, E.J., Ziolkowska, L., Griffin, C., Choi, K., Franklin, F.C.H., Martienssen, R.A., et al. (2017) Natural variation and dosage of the HEI10 meiotic E3 ligase control Arabidopsis crossover recombination. Genes Dev, 31, 306–317.

8. Morgan, C., Fozard, J.A., Hartley, M., Henderson, I.R., Bomblies, K. and Howard, M. (2021) Diffusion-mediated HEI10 coarsening can explain meiotic crossover positioning in Arabidopsis. Nat Commun, 12.

9. Lian, Q., Jing, J., Ernst, M., Grelon, M. and Mercier, R. (2022) Joint control of meiotic crossover patterning by the synaptonemal complex and HEI10 dosage. Nature Commun, 10.1038/s41467-022-33472-w.

10. Morgan, C., Howard, M. and Henderson, I.R. (2024) HEI10 coarsening, chromatin and sequence polymorphism shape the plant meiotic recombination landscape. Curr Opin Plant Biol, 81, 102570.

11. Cannavo, E., Sanchez, A., Anand, R., Ranjha, L., Hugener, J., Adam, C., Acharya, A., Weyland, N., Aran-Guiu, X., Charbonnier, J.-B., et al. (2020) Regulation of the MLH1-MLH3 endonuclease in meiosis. Nature, 10.1101/2020.02.12.946293.

12. Kulkarni, D., Owens, S., Honda, M., Ito, M., Yang, Y., Corrigan, M., Chen, L., Quan, A. and Hunter, N. (2020) PCNA activates the MutLγ endonuclease to promote meiotic crossing over. Nature, 586, 623–627.

13. Barlow, A.L. and Hultén, M.A. (1998) Crossing over analysis at pachytene in man. Eur J Hum Genet, 6, 350–358.

14. Anderson, L.K., Reeves, A., Webb, L.M. and Ashley, T. (1999) Distribution of crossing over on mouse synaptonemal complexes using immunofluorescent localization of MLH1 protein. Genetics, 151, 1569–1579.

15. Jackson, N., Sanchez-Moran, E., Buckling, E., Armstrong, S.J., Jones, G.H. and Franklin, F.C.H. (2006) Reduced meiotic crossovers and delayed prophase I progression in AtMLH3-deficient Arabidopsis. EMBO J, 25, 1315–23.

16. Dion, E., Li, L., Jean, M. and Belzile, F. (2007) An Arabidopsis MLH1 mutant exhibits reproductive defects and reveals a dual role for this gene in mitotic recombination. Plant J, 51, 431–40.

17. Underwood, C.J., Choi, K., Lambing, C., Zhao, X., Serra, H., Borges, F., Simorowski, J., Ernst, E., Jacob, Y., Henderson, I.R., et al. (2018) Epigenetic activation of meiotic recombination near Arabidopsis thaliana centromeres via loss of H3K9me2 and non-CG DNA methylation. Genome Res, 28, 519–531.

18. Alou, A.H., Jean, M., Domingue, O. and Belzile, F.J. (2004) Structure and expression of AtPMS1, the Arabidopsis ortholog of the yeast DNA repair gene PMS1. Plant Science, 167, 447–456.

19. Wang, T.F., Kleckner, N. and Hunter, N. (1999) Functional specificity of MutL homologs in yeast: evidence for three Mlh1-based heterocomplexes with distinct roles during meiosis in recombination and mismatch correction. Proc Natl Acad Sci U S A, 96, 13914–13919.

20. Li, L., Dion, E., Richard, G., Domingue, O., Jean, M. and Belzile, F.J. (2009) The Arabidopsis DNA mismatch repair gene PMS1 restricts somatic recombination between homeologous sequences. Plant Mol Biol, 69, 675–84.

21. Higgins, J.D., Buckling, E.F., Franklin, F.C.H. and Jones, G.H. (2008) Expression and functional analysis of AtMUS81 in Arabidopsis meiosis reveals a role in the second pathway of crossing-over. Plant J, 54, 152–62.

22. Berchowitz, L.E., Francis, K.E., Bey, A.L. and Copenhaver, G.P. (2007) The Role of AtMUS81 in Interference-Insensitive Crossovers in A. thaliana. PLoS Genet, 3, e132.

23. Kurzbauer, M.T., Pradillo, M., Kerzendorfer, C., Sims, J., Ladurner, R., Oliver, C., Janisiw, M.P., Mosiolek, M., Schweizer, D., Copenhaver, G.P., et al. (2018) Arabidopsis thaliana FANCD2 promotes meiotic crossover formation. Plant Cell, 30, 415–428.

24. Li, X., Zhang, J., Huang, J., Xu, J., Chen, Z., Copenhaver, G.P. and Wang, Y. (2021) Regulation of interference-sensitive crossover distribution ensures crossover assurance in Arabidopsis. Proc Natl Acad Sci U S A, 118, e2107543118.

25. Hartung, F., Suer, S. and Puchta, H. (2007) Two closely related RecQ helicases have antagonistic roles in homologous recombination and DNA repair in Arabidopsis thaliana. Proc Natl Acad Sci U S A, 104, 18836–18841.

26. Girard, C., Chelysheva, L., Choinard, S., Froger, N., Macaisne, N., Lehmemdi, A., Mazel, J., Crismani, W. and Mercier, R. (2015) AAA-ATPase FIDGETIN-LIKE 1 and Helicase FANCM Antagonize Meiotic Crossovers by Distinct Mechanisms. PLoS Genet, 11, e1005369.

27. Séguéla-Arnaud, M., Crismani, W., Larchevêque, C., Mazel, J., Froger, N., Choinard, S., Lemhemdi, A., Macaisne, N., Van Leene, J., Gevaert, K., et al. (2015) Multiple mechanisms limit meiotic crossovers: TOP3α and two BLM homologs antagonize crossovers in parallel to FANCM. Proceedings of the National Academy of Sciences, 112, 4713–4718.

28. Crismani, W., Girard, C., Froger, N., Pradillo, M., Santos, J.L., Chelysheva, L., Copenhaver, G.P., Horlow, C. and Mercier, R. (2012) FANCM limits meiotic crossovers. Science, 336, 1588– 1590.

29. Melamed-Bessudo, C., Yehuda, E., Stuitje, A.R. and Levy, A. a (2005) A new seed-based assay for meiotic recombination in Arabidopsis thaliana. The Plant Journal, 43, 458– 466.

30. Hord, C.L.H., Sun, Y.J., Pillitteri, L.J., Torii, K.U., Wang, H., Zhang, S. and Ma, H. (2008) Regulation of Arabidopsis early anther development by the mitogen-activated protein kinases, MPK3 and MPK6, and the ERECTA and related receptor-like kinases. Mol Plant, 1, 645–658.

31. Alexander, M.P. (1969) Differential staining of aborted and nonaborted pollen. Biotechnic and Histochemistry, 44, 117–122.

32. Wu, G., Rossidivito, G., Hu, T., Berlyand, Y. and Poethig, R.S. (2015) Traffic lines: New tools for genetic analysis in Arabidopsis thaliana. Genetics, 200, 35–45.

33. Kbiri, N., Dluzewska, J., Henderson, I.R. and Ziolkowski, P.A. (2022) Quantifying Meiotic Crossover Recombination in Arabidopsis Lines Expressing Fluorescent Reporters in Seeds Using SeedScoring Pipeline for CellProfiler. In Methods in Molecular Biology. Humana Press Inc., Vol. 2484, pp. 121–134.

34. Martinez-Garcia, M. and Pradillo, M. (2017) Functional analysis of Arabidopsis argonautes in meiosis and DNA repair. In Methods in Molecular Biology. Humana Press Inc., Vol. 1640, pp. 145–158.

35. Zhu, L., Fernández-Jiménez, N., Szymanska-Lejman, M., Pelé, A., Underwood, C.J., Serra, H., Lambing, C., Dluzewska, J., Bieluszewski, T., Pradillo, M., et al. (2021) Natural variation identifies SNI1, the SMC5/6 component, as a modifier of meiotic crossover in Arabidopsis. Proceedings of the National Academy of Sciences, 118, e2021970118.

36. Sanchez Moran, E., Armstrong, S.J., Santos, J.L., Franklin, F.C.H. and Jones, G.H. (2001) Chiasma formation in Arabidopsis thaliana accession Wassileskija and in two meiotic mutants. Chromosome Research, 9, 121–128 (2001).

37. Ross, K.J., Fransz, P. and Jones, G.H. (1996) A light microscopic atlas of meiosis in Arabidopsis thaliana. Chromosome Research, 4, 507–516.

38. Gerlach, W.L. and Bedbrook, J.R. (1979) Cloning and characterization of ribosomal RNA genes from wheat and barley. Nucleic Acids Res, 7.

39. Campell, B.R., Song, Y., Posch, T.E., Cullis, C.A. and Town, C.D. (1992) Sequence and organization of 5S ribosomal RNA-encoding genes of Arabidopsis thaliana. Gene, 112, 225–228.

40. Bieluszewski, T., Szymanska-Lejman, M., Dziegielewski, W., Zhu, L. and Ziolkowski, P.A. (2022) Efficient Generation of CRISPR/Cas9-Based Mutants Supported by Fluorescent Seed Selection in Different Arabidopsis Accessions. In Methods in Molecular Biology. Humana Press Inc., Vol. 2484, pp. 161–182.

41. Rowan, B.A., Patel, V., Weigel, D. and Schneeberger, K. (2015) Rapid and Inexpensive Whole-Genome Genotyping-by-Sequencing for Crossover Localization and Fine-Scale Genetic Mapping. G3: Genes|Genomes|Genetics, 5, 385–398.

42. Szymanska-Lejman, M., Dziegielewski, W., Dluzewska, J., Kbiri, N., Bieluszewska, A., Poethig, R.S. and Ziolkowski, P.A. (2023) The effect of DNA polymorphisms and natural variation on crossover hotspot activity in Arabidopsis hybrids. Nat Commun, 14, 33.

43. Dluzewska, J., Dziegielewski, W., Szymanska-Lejman, M., Gazecka, M., Henderson, I.R., Higgins, J.D. and Ziolkowski, P.A. (2023) MSH2 stimulates interfering and inhibits non-interfering crossovers in response to genetic polymorphism. Nat Commun, 14, 6716.

44. Langmead, B. and Salzberg, S.L. (2012) Fast gapped-read alignment with Bowtie 2. Nat Methods, 9, 357–359.

45. Li, H., Handsaker, B., Wysoker, A., Fennell, T., Ruan, J., Homer, N., Marth, G., Abecasis, G. and Durbin, R. (2009) The Sequence Alignment/Map format and SAMtools. Bioinformatics, 25, 2078–2079.

46. Li, H. (2011) A statistical framework for SNP calling, mutation discovery, association mapping and population genetical parameter estimation from sequencing data. Bioinformatics, 27, 2987–2993.

47. Rowan, B.A., Heavens, D., Feuerborn, T.R., Tock, A.J., Henderson, I.R. and Weigel, D. (2019) An Ultra High-Density Arabidopsis thaliana Crossover. Genetics, 213, 771–787.

48. Langmea, B. and Salzberg, S.L. (2013) Fast gapped-read alignment with Bowtie 2. Nat Methods, 9, 357–359.

49. Li, H. and Durbin, R. (2009) Fast and accurate short read alignment with Burrows-Wheeler transform. Bioinformatics, 25, 1754–1760.

50. Pedersen, B.S. and Quinlan, A.R. (2018) Mosdepth: Quick coverage calculation for genomes and exomes. Bioinformatics, 34, 867–868.

51. Wang, Y. and Copenhaver, G.P. (2018) Meiotic Recombination: Mixing It Up in Plants. Annu Rev Plant Biol, 69, 577–609.

52. Ziolkowski, P.A. (2022) Why do plants need the ZMM crossover pathway? A snapshot of meiotic recombination from the perspective of interhomolog polymorphism. Plant Reprod, 1, 3.

53. Mercier, R., Mézard, C., Jenczewski, E., Macaisne, N. and Grelon, M. (2015) The Molecular Biology of Meiosis in Plants. Annu Rev Plant Biol, 66, 297–327.

54. Henry, I.M., Dilkes, B.P., Miller, E.S., Burkart-Waco, D. and Comai, L. (2010) Phenotypic consequences of aneuploidy in Arabidopsis thaliana. Genetics, 186, 1231–1245.

55. Serra, H., Lambing, C., Griffin, C.H., Topp, S.D., Nageswaran, D.C., Underwood, C.J., Ziolkowski, P.A., Séguéla-Arnaud, M., Fernandes, J.B., Mercier, R., et al. (2018) Massive crossover elevation via combination of HEI10 and recq4a recq4b during Arabidopsis meiosis. Proc Natl Acad Sci U S A, 115, 2437–2442.

56. Blackwell, A.R., Dluzewska, J., Szymanska-Lejman, M., Desjardins, S., Tock, A.J., Kbiri, N., Lambing, C., Lawrence, E.J., Bieluszewski, T., Rowan, B., et al. (2020) MSH2 shapes the meiotic crossover landscape in relation to interhomolog polymorphism in Arabidopsis. EMBO Journal, 39, e104858.

57. Drouaud, J., Camilleri, C., Bourguignon, P., Canaguier, A., Bérard, A., Vezon, D., Giancola, S., Brunel, D., Colot, V., Prum, B., et al. (2006) Variation in crossing-over rates across chromosome 4 of Arabidopsis thaliana reveals the presence of meiotic recombination “hot spots”. Genome Res, 16, 106–114.

58. Sun, H., Rowan, B.A., Flood, P.J., Brandt, R., Fuss, J., Hancock, A.M., Michelmore, R.W., Huettel, B. and Schneeberger, K. (2019) Linked-read sequencing of gametes allows efficient genome-wide analysis of meiotic recombination. Nat Commun, 10, 4310.

59. Chelysheva, L., Gendrot, G., Vezon, D., Doutriaux, M.-P., Mercier, R. and Grelon, M. (2007) Zip4/Spo22 is required for class I CO formation but not for synapsis completion in Arabidopsis thaliana. PLoS Genet, 3, e83.

60. Knoll, A., Higgins, J.D., Seeliger, K., Reha, S.J., Dangel, N.J., Bauknecht, M., Schröpfer, S., Franklin, F.C.H. and Puchta, H. (2012) The Fanconi anemia ortholog FANCM ensures ordered homologous recombination in both somatic and meiotic cells in Arabidopsis. Plant Cell, 24, 1448–64.

61. Lloyd, A. (2022) Crossover patterning in plants. Plant Reprod, 10.1007/s00497-022-00445-4.

62. Al-Sweel, N., Raghavan, V., Dutta, A., Ajith, V.P., Di Vietro, L., Khondakar, N., Manhart, C.M., Surtees, J.A., Nishant, K.T. and Alani, E. (2017) mlh3 mutations in baker’s yeast alter meiotic recombination outcomes by increasing noncrossover events genome-wide. PLoS Genet, 13, e1006974.

63. Claeys Bouuaert, C. and Keeney, S. (2017) Distinct DNA-binding surfaces in the ATPase and linker domains of MutLγ determine its substrate specificities and exert separable functions in meiotic recombination and mismatch repair. PLoS Genet, 13, e1006722.

64. Pyatnitskaya, A., Andreani, J., Guérois, R., De Muyt, A. and Borde, V. (2022) The Zip4 protein directly couples meiotic crossover formation to synaptonemal complex assembly. Genes Dev, 36, 53–63.

65. Zhang, Q., Ji, S.Y., Busayavalasa, K. and Yu, C. (2019) SPO16 binds SHOC1 to promote homologous recombination and crossing-over in meiotic prophase I. Sci Adv, 5.

66. Toledo, M., Sun, X., Brieño-Enríquez, M.A., Raghavan, V., Gray, S., Pea, J., Milano, C.R., Venkatesh, A., Patel, L., Borst, P.L., et al. (2019) A mutation in the endonuclease domain of mouse MLH3 reveals novel roles for mutlγ during crossover formation in meiotic prophase I. PLoS Genet, 15.

67. Lipkin, S.M., Moens, P.B., Wang, V., Lenzi, M., Shanmugarajah, D., Gilgeous, A., Thomas, J., Cheng, J., Touchman, J.W., Green, E.D., et al. (2002) Meiotic arrest and aneuploidy in MLH3-deficient mice. Nat Genet, 31, 385–390.

68. Osman, K., Sanchez-Moran, E., Mann, S.C., Jones, G.H. and Franklin, F.C.H. (2009) Replication protein A (AtRPA1a) is required for class I crossover formation but is dispensable for meiotic DNA break repair. EMBO J, 28, 394–404.

69. Chelysheva, L., Vezon, D., Chambon, A., Gendrot, G., Pereira, L., Lemhemdi, A., Vrielynck, N., Le Guin, S., Novatchkova, M. and Grelon, M. (2012) The Arabidopsis HEI10 is a new ZMM protein related to Zip3. PLoS Genet, 8, e1002799.

70. Blackwell, A.R., Dluzewska, J., Szymanska-Lejman, M., Desjardins, S., Tock, A.J., Kbiri, N., Lambing, C., Lawrence, E.J., Bieluszewski, T., Rowan, B., et al. (2020) MSH2 shapes the meiotic crossover landscape in relation to interhomolog polymorphism in Arabidopsis. EMBO Journal, 39, e104858.

71. Blary, A., Gonzalo, A., Eber, F., Bérard, A., Bergès, H., Bessoltane, N., Charif, D., Charpentier, C., Cromer, L., Fourment, J., et al. (2018) FANCM limits meiotic crossovers in Brassica crops. Front Plant Sci, 368, 9.

72. Desjardins, S.D., Simmonds, J., Guterman, I., Kanyuka, K., Burridge, A.J., Tock, A.J., Sanchez-moran, E., Franklin, F.C.H., Henderson, I.R., Edwards, K.J., et al. (2022) FANCM promotes class I interfering crossovers and suppresses class II non-interfering crossovers in wheat meiosis. Nat Commun, 3, 3644.

73. Mieulet, D., Aubert, G., Bres, C., Klein, A., Droc, gaetan G., Vieille, E., Rond-Coissieux, C., Sanchez, Myriam., Dalmais, M., Mauxion, P., et al. (2018) Unleashing meiotic crossovers in crops. Nat Plants, 10.1073/pnas.92.19.8675.

74. De Muyt, A., Jessop, L., Kolar, E., Sourirajan, A., Chen, J., Dayani, Y. and Lichten, M. (2012) BLM Helicase Ortholog Sgs1 Is a Central Regulator of Meiotic Recombination Intermediate Metabolism. Mol Cell, 46, 43–53.

75. Zakharyevich, K., Tang, S., Ma, Y. and Hunter, N. (2012) Delineation of joint molecule resolution pathways in meiosis identifies a crossover-specific resolvase. Cell, 149, 334– 347.

76. Pannafino, G., Chen, J.J., Mithani, V., Payero, L., Gioia, M., Crickard, J.B. and Alani, E. (2024) The Dmc1 recombinase physically interacts with and promotes the meiotic crossover functions of the Mlh1-Mlh3 endonuclease. Genetics, 227, 1–15.

