## Supplementary Figures for "Genetic dissection of MutL complexes in Arabidopsis meiosis"

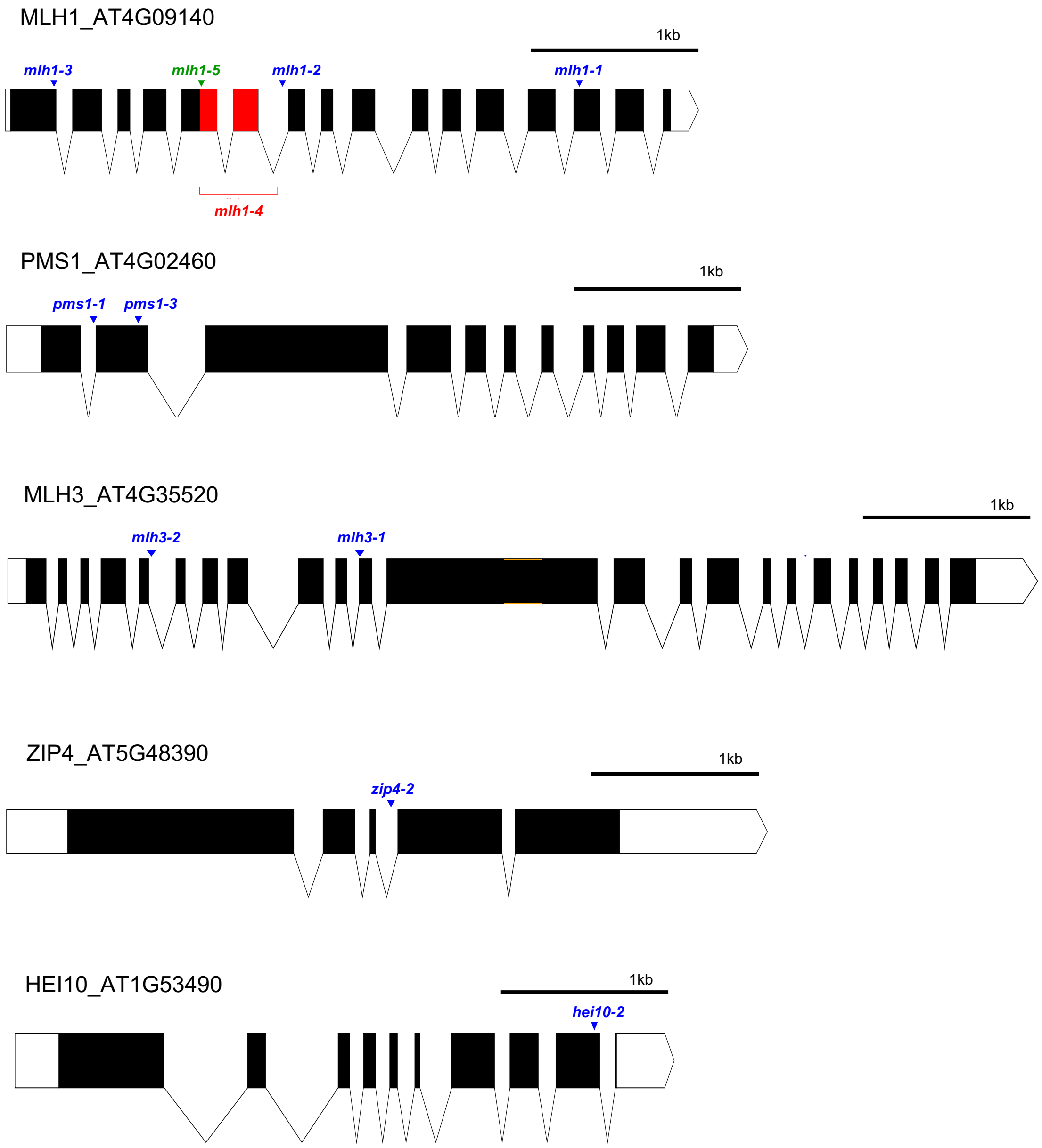


**Supplementary Figure S1. Scaled representation of the genomic sequences of the MutL genes *MLH1, PMS1,* and *MLH3* and the ZMM genes *ZIP4* and *HEI10*.** 3' and 5' UTRs are represented by white blocks, exons by black blocks, and introns by V-shaped linkers. T-DNA insertion mutations are represented by blue arrowheads and text, the SNP mutation is represented by a green arrowhead and text, and the deletion mutation is represented by a red bracket, text, and deleted exons. AGI codes are provided. Scale bar, 1kb.

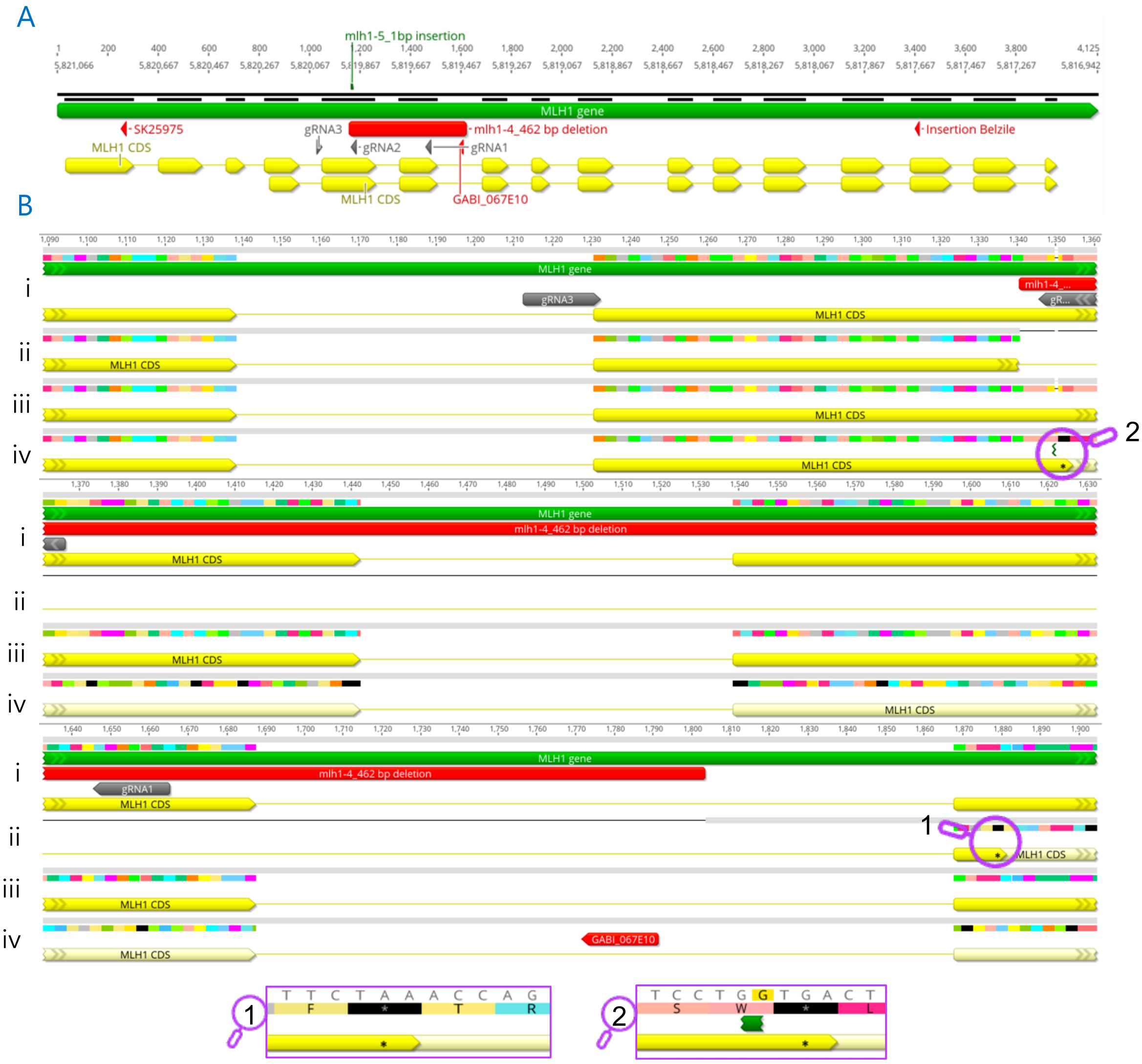


**Supplementary Figure S2.** ***mlh1-4* and *mlh1-5* CRISPR-Cas9 mediated mutants in Col-0 and L*er*-0 respectively. A**, *MLH1* gene structure. The exons are represented with yellow arrows, the gRNAs used for the mutagenesis in grey arrowheads, the obtained 462 bp deletion in Col-0 is represented with a red rectangle, and the 1 bp insertion in Ler-0 is represented with a green flag. The position of the T-DNA insertion of the other MLH1 mutants is also represented with red arrowheads. The genomic position is represented with the scale. **B**, *mlh1-4* and *mlh1-5* are aligned to the wildtype references of Col-0 and L*er*-0. **i.** Col-0 WT*.* **ii.** *mlh1-4* allele. **iii.** L*er*-0 WT. iv. *mlh1-5* allele*.* The 462 bp genomic and 250 bp in coding sequence deletion, in Col-0, introduces a frameshift and multiple STOP codons. Similarly, the 1bp insertion in L*er*-0 also introduces a frameshift and multiple STOP codons. STOP codons are represented in black, the predicted first STOP codons and the predicted end of the transcripts, are highlighted with purple magnifying glasses: **1** for *mlh1-4* and **2** for *mlh1-5*. The reverse sequence is used for simplicity.

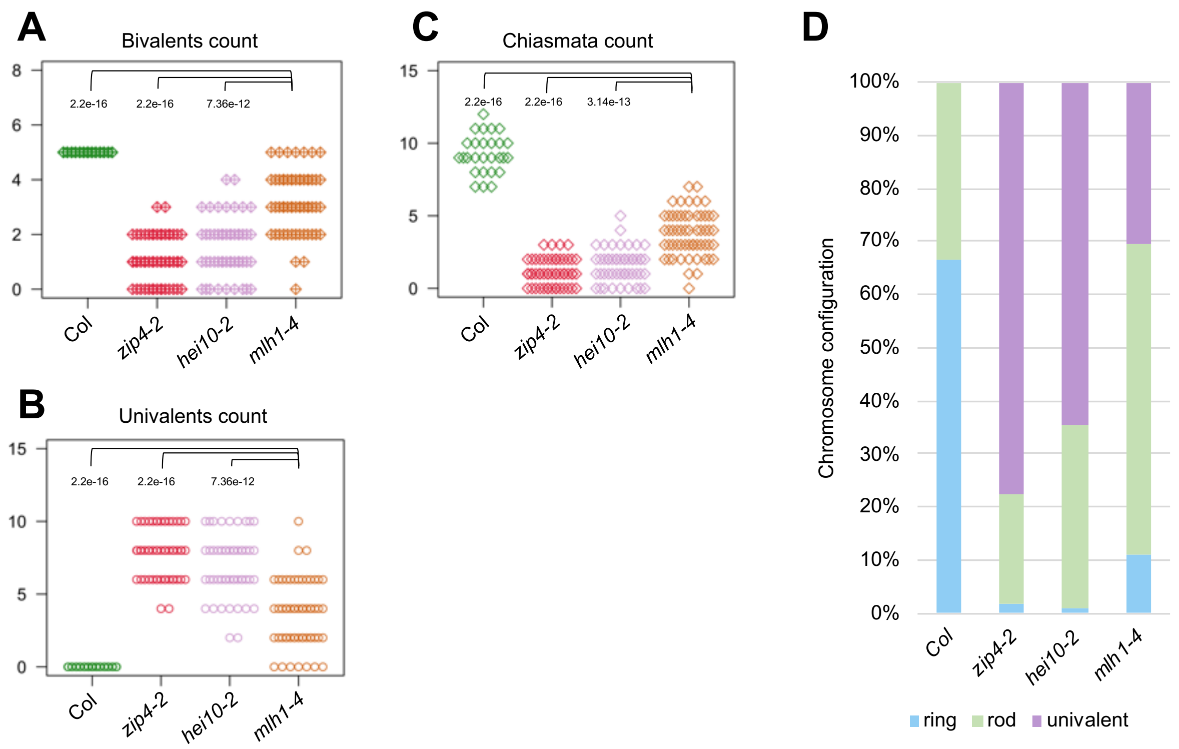


**Supplementary Figure S3. Comparative analysis of the meiotic behavior of Col, *zip4-2, hei10-2,* and *mlh1-4.* A-C.** Graphical representation of bivalent count per metaphase I meiocyte (**A**), univalent count (**B**), and chiasmata count (**C**). The *P* values were estimated using a Welch t-test. **D.** Chromosome configuration represented in ring, rod, and univalent proportions. A summary table of the average numbers of univalents, bivalents, and chiasmata is provided in Supplementary Table S6.

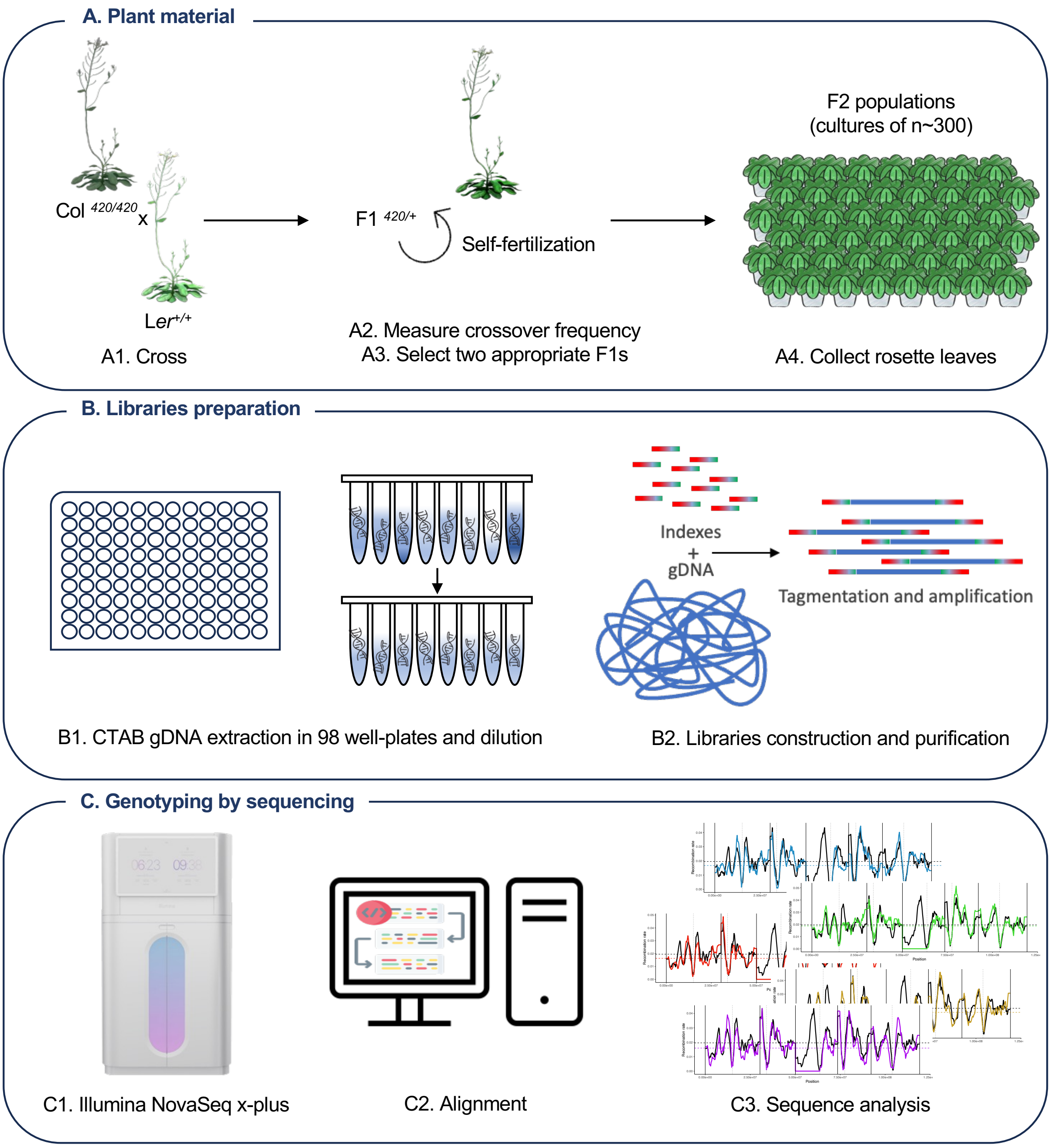


**Supplementary Figure S4. Generation of genome-wide crossover maps. A.** Plant material preparation, the Col$^{420/420}$ and L*er*$^{+/+}$ Arabidopsis genetic background were chosen to generate the appropriate mutants and then crossed to create an $F_1^{420/+}$ population (**A1**). The $F_1^{420/+}$ were grown to seed, crossover recombination frequency was measured in the 420 interval (**A2**), and 2 individuals were selected based on having an RF closest to the average of the population and the best fluorescent tags segregation ratios (**A3**). Next, about 300 seeds were sown to construct an $F_2$ population and rosette leaves were collected for gDNA extraction. **B.** gDNA libraries were constructed by extracting gDNA from rosette leaves using CTAB protocol. The DNA samples were diluted to a concentration of 5ng/µL (**B1**), then tagmented using Tn5 and amplified with KAPA2G Robust to introduce indexes. The pooled, size selected (450bp-

700bp) and purified libraries were then sent for sequencing (**B2**). **C.** gDNA libraries were sequenced using NovaSeq x-plus and demultiplexed by Macrogen Europe (**C1**), the received data was aligned to a Col reference and computed (**C2**) to allow for different types of sequence analyses (**C3**). The detailed computation of genome-wide sequencing is provided in materials and methods. Rights to the Arabidopsis plant drawing belong to @_HETAKA, https://doi.org/10.7875/togopic.2021.057, the plant in a pot was designed by Freepik, and the Illumina NovaSeq x-plus was taken from Illumina's official website https://emea.illumina.com.

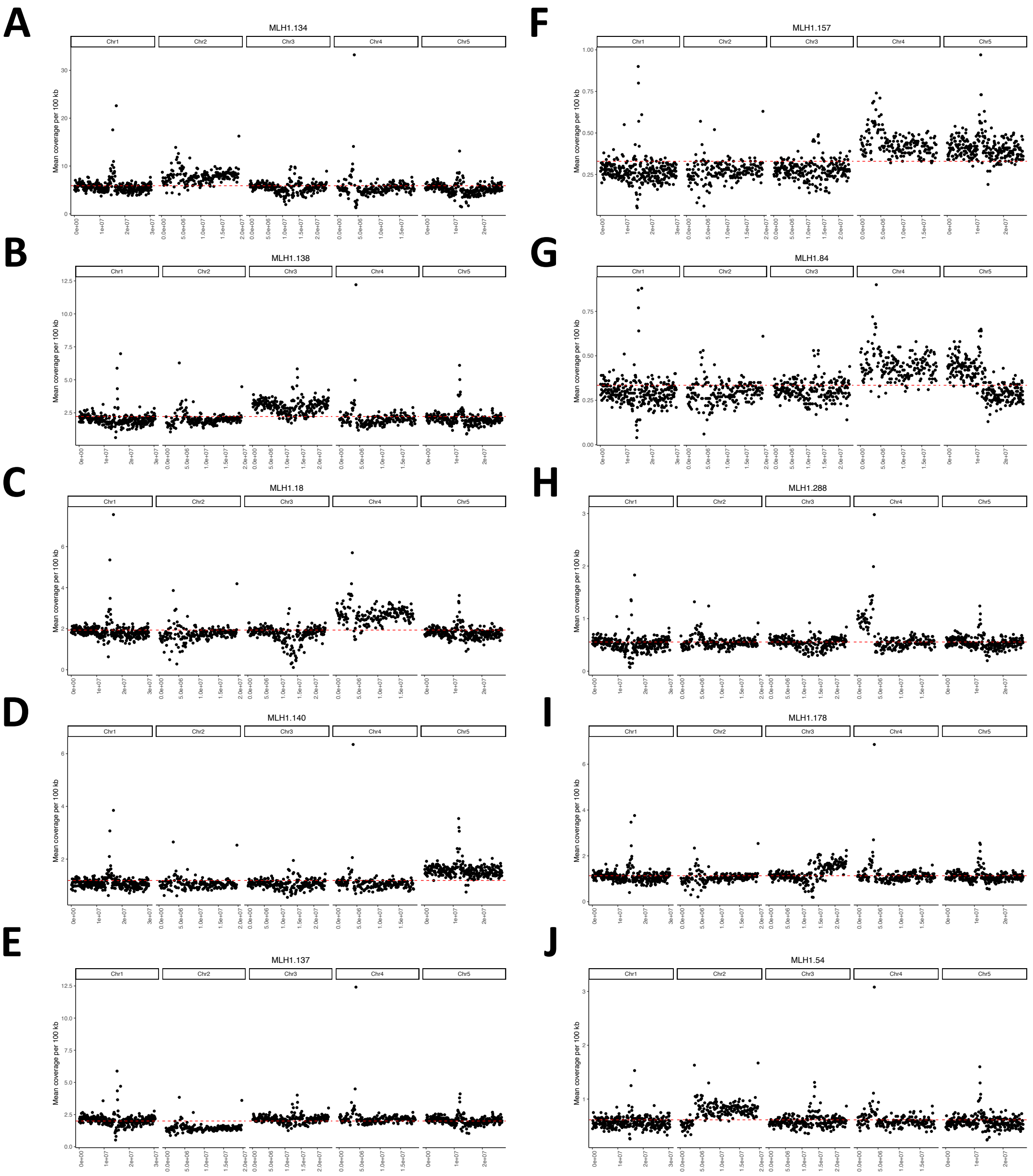


**Supplementary Figure S5. Examples of all types of aneuploidy found in Col/L*er mlh1* $F_2$ individuals.** Mean sequence coverage plots per 100 kb. **A.** Trisomy of chromosome 2. **B.** Trisomy of chromosome 3 . **C.** Trisomy of chromosome 4. **D.** Trisomy of chromosome 5. **E.** Monosomy of chromosome 2. **F.** Trisomy of chromosomes 4 and 5. **G.** Trisomy of chromosome 4 and north arm of chromosome 5. **H.** Partial trisomy of north arm of chromosome 4. **I.** Partial trisomy of south arm of chromosome 3. **J.** Partial trisomy of south arm of chromosome 2.

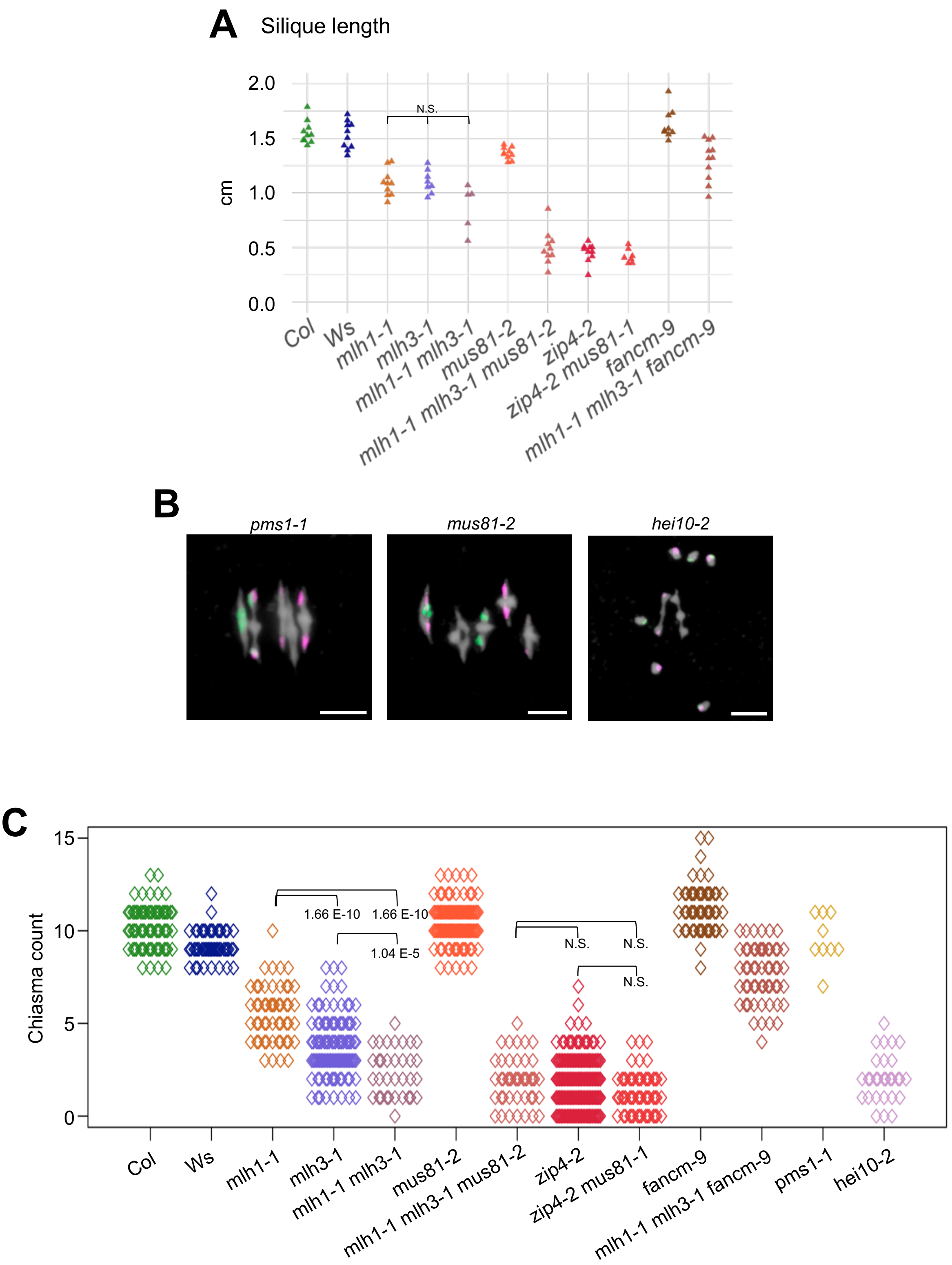


**Supplementary Figure S6. Additional information for characterizing the genetic interaction between MutLγ loss of function and class II crossover recombination loss of function *mus81* and *fancm*. A.** Fertility assessed by silique length. The phenotype is consistent with the seed set and pollen viability in Figure 6. The *P* values were estimated using one-way ANOVA and the Tukey HSD tests (Supplementary Tables S7-S8). n = 5 to 11. **B.**

Cytological characterization at metaphase I for *pms1, mus81*, and *hei10.* The *pms1* and *mus81* mutations do not affect bivalent formation, whereas *hei10,* as most *zmm* mutants, shows dramatically reduced chiasmata/ bivalents formation. **C.** Chiasmata count per meiocyte for all tested lines. The observed phenotype is consistent with the fertility assessment and cytological analysis. The *P* values were estimated using one-way ANOVA and the Tukey HSD/Tukey-Kramer tests (Supplementary Table S16). The number of characterized meiocytes are Col (n = 69), Ws (n = 50)*, mlh1* (n = 53)*, mlh3* (n = 108), *mlh1 mlh3* (n = 34), *mus81* (n = 102), *mlh1 mlh3 mus81* (n = 44), *zip4* (n=255), *zip4 mus81* (n=57), *fancm* (n = 52), *mlh1 mlh3 fancm* (n = 55), *pms1* (n= 9) and *hei10* (n= 29).

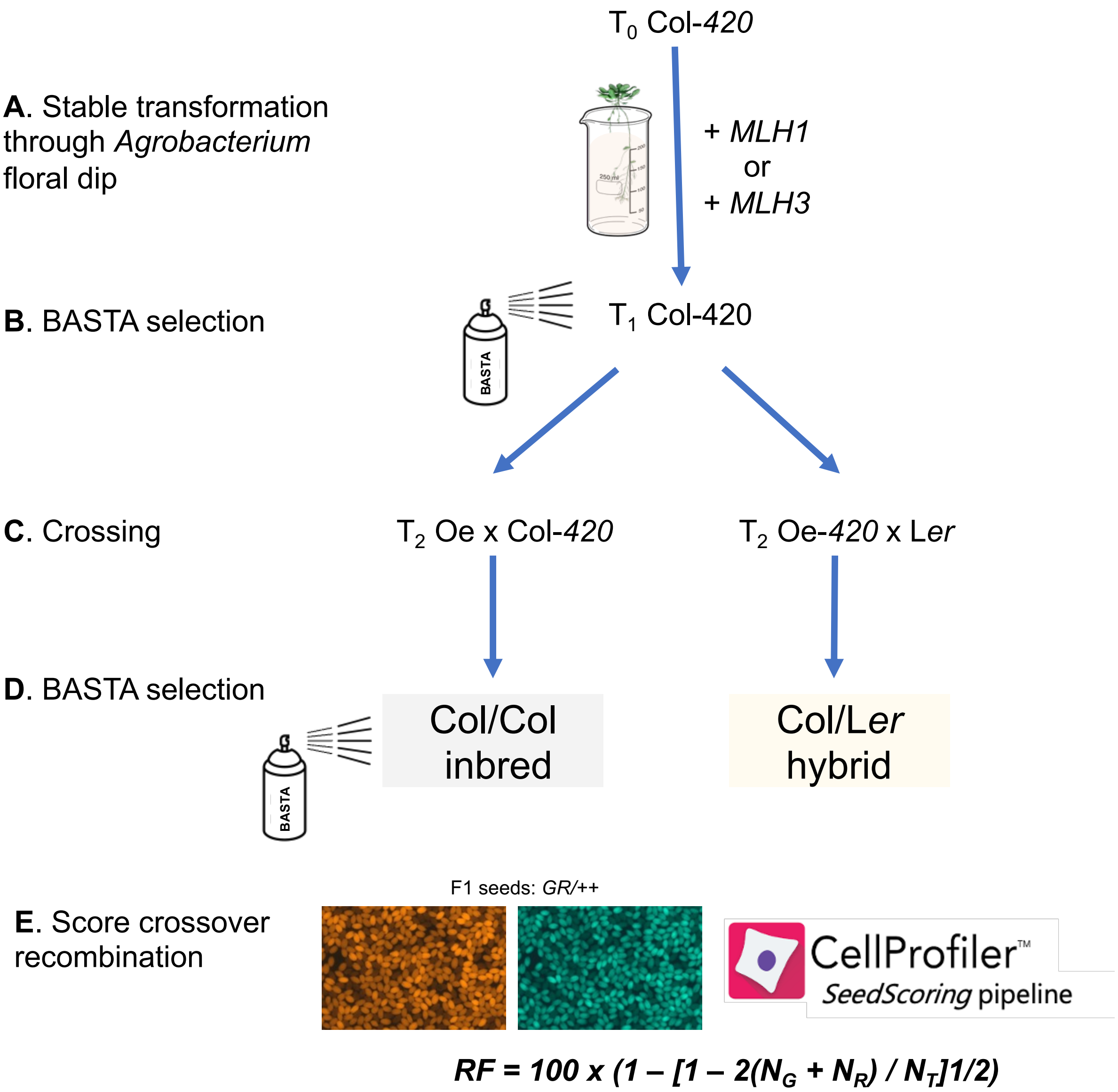


**Supplementary Figure S7. Experimental design for measuring crossover frequency in Col/Col inbred and Col/*Ler* hybrid contexts, for MutL overexpressors. A.** $T_0$ Col$^{420/+}$ plants were transformed with constructs harboring genomic sequences of *MLH1* or *MLH3* under their respective native promoters or the meiosis-specific DMC1 promoter. They were grown to seed and collected. **B.** $T_1$ seeds were sown and $T_1$ plants were selected for transformants using BASTA. $T_1$ transformants were grown to seed and collected. **C.** $T_2$ seeds were sown and $T_2$ plants were selected with BASTA then crossed to Col and L*er*, F1 seeds were collected. **D.** $F_1$ plants were grown, selected with BASTA, and collected. **E.** The obtained $F_2$ seeds were pictured following the protocol in (6) to measure crossover recombination frequency in the *420* interval.

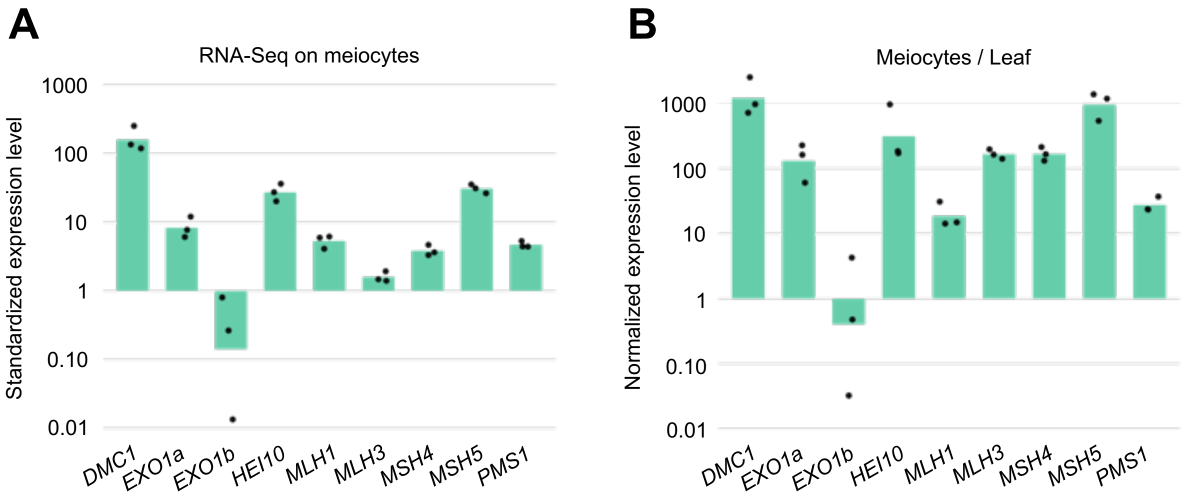


**Supplemental Figure S8. Physiological expression levels of selected genes in wildtype Arabidopsis. A.** Standardized expression levels of *DMC1, EXO1a, EXO1b, HEI10, MLH1, MLH3, MSH4, MSH5,* and *PMS1* in meiocytes. **B.** Meiocyte to leaf normalized expression level of the same genes. n=3. Open access data from Walker et al.,2017.

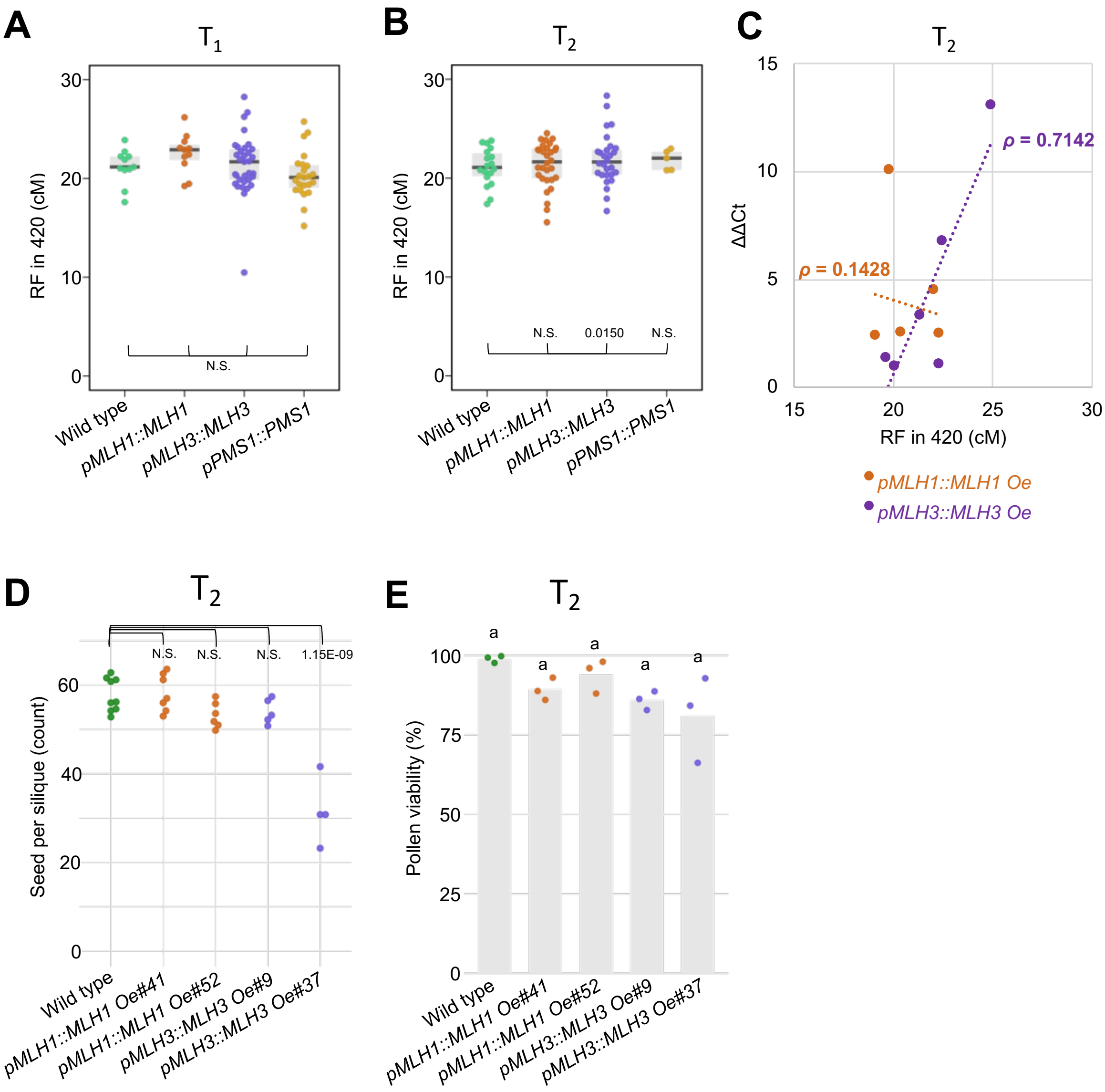


**Supplementary Figure S9. Additional information on the phenotypical characterization of *MutL* overexpressors under the control of their respective native promoters. A-B.** Local meiotic crossover recombination frequency, measured in the *420* interval, for *MLH1, MLH3,* and *PMS1* in $T_1$ generation (**A**), and $T_2$ generation (**B**). The *P* values were estimated using the Welch t-test. **C.** Crossover recombination frequency correlates positively with *MLH3* expression level, Spearman *Rho* = 0.7142, whereas *MLH1* expression level does not, Spearman *Rho* = 0.1428. n= 3 x 3 biological and technical replicates. **D-E.** Fertility assays for *MLH1* and *MLH3* overexpressors were assessed by seed set (**D**), and pollen viability (**E**). The *P* values were estimated using one-way ANOVA and the Tukey HSD tests (Supplementary Tables S19-S20). n = 4 to 9 in D and n = 3 in E. Expression level quantification using RT-qPCR and fertility assays were performed on $T_2$ generation plants.

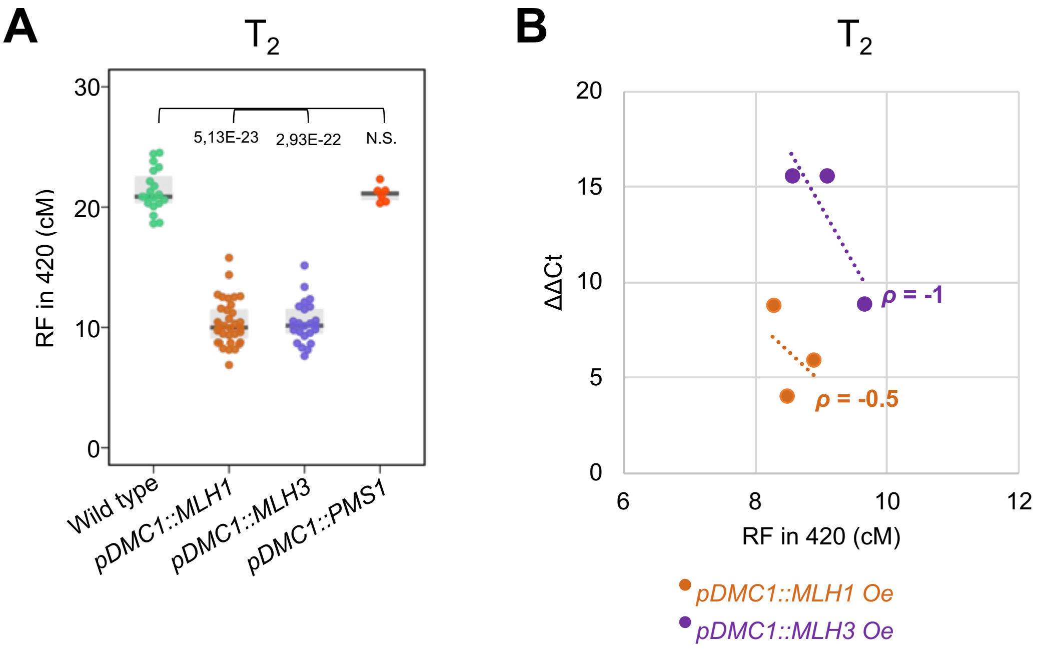


**Supplementary Figure S10. Additional information on the phenotypical characterization of *MutL* overexpressors under the control of *DMC1* promoter. A.** Local meiotic crossover recombination frequency, measured in the *420* interval, for *MLH1, MLH3,* and *PMS1* under *DMC1* promoter in $T_2$ generation. The *P* values were estimated using the Welch t-test. **B.** Crossover recombination frequency correlates negatively with *MLH1* and *MLH3* expression levels, Spearman *Rho* = -0.5 and -1, respectively. n = 3 x 3 biological and technical replicates for *MLH3* and n= 1 x 3 for *MLH1*.

**Supplementary Tables (in Excel file)**

**Supplementary Table S1.** Genotyping primers used for discriminating the different used mutants.

**Supplementary Table S2.** CRISPR Cas9 gRNA and cloning primer sequences used for knocking out *MLH1.*

**Supplementary Table S3.** Primer sequences used for cloning MLH1, MLH3 and PMS1 under their respective native promoters or inframe with the DMC1 promoter for ectopic expression.

**Supplementary Table S4.** Summary of the sequencing data for Col/L*er mlh1* and *hei10-2/+.*

**Supplementary Table S5.** The qPCR primer sequences used to quantify the different specified targets following the reverse transcription of total RNA.

**Supplementary Table S6.** Summary table of the average numbers of univalents, bivalents, and chiasmata in the tested lines.

**Supplementary Table S7.** The average number of chiasmata per chromosome and Pollen Mother Cells (PMC) of MutL mutants in combination with class II factors at Metaphase I.

**Supplementary Table S8.** Meiotic behavior established from Pollen Mother Cells of MutL mutants in combination with class II factors at Metaphase I.

**Supplementary Table S9.** Chromosome configuration proportions in all tested lines. Submetacentric (1+3+5) and acrocentric (2+4) chromosomes behavior is discriminated.

**Supplementary Table S10.** Multidirectional One way ANOVA Tukey HSD test on seed set data in Figure 1b.

**Supplementary Table S11.** Multidirectional One way ANOVA Tukey HSD test on silique length data in Figure 1c.

**Supplementary Table S12.** Multidirectional One way ANOVA Tukey HSD test on pollen viability in Figure 1d.

**Supplementary Table S13.** Multidirectional One way ANOVA Tukey HSD test on seed set data in Figure 6a.

**Supplementary Table S14.** Multidirectional One way ANOVA Tukey HSD test on pollen viability data in Figure 6b.

**Supplementary Table S15.** Multidirectional One way ANOVA Tukey HSD test on silique length data in Supplementary Figure 6a.

**Supplementary Table S16.** Multidirectional One way ANOVA Tukey HSD test on chiasmata count data in Supplementary Figure 6c.

**Supplementary Table S17.** Multidirectional One way ANOVA Tukey HSD test on seed set data in Figure 7d.

**Supplementary Table S18.** Multidirectional One way ANOVA Tukey HSD test on pollen viability data in Figure 7e.

**Supplementary Table S19.** Multidirectional One way ANOVA Tukey HSD test of seed set data in Supplementary Figure 9d.

**Supplementary Table S20.** Multidirectional One way ANOVA Tukey HSD test on pollen viability data in Supplementary Figure 9e.
